# A Subcortical Feeding Circuit Linking Interoception to Jaw movement

**DOI:** 10.1101/2023.10.11.561735

**Authors:** Christin Kosse, Jessica Ivanov, Zachary Knight, Kyle Pellegrino, Jeffrey Friedman

## Abstract

The brain processes an array of stimuli enabling the selection of an appropriate behavioural response but the neural pathways linking interoceptive inputs to outputs for feeding are poorly understood. Here we delineate a subcortical circuit in which brain-derived neurotrophic factor (BDNF) expressing neurons in the ventromedial hypothalamus (VMH) directly connect interoceptive inputs to motor centers controlling food consumption and jaw movements. VMH^BDNF^ neuron inhibition increases food intake by gating motor sequences of feeding through projections to premotor areas of the jaw. When food is unavailable, VMH^BDNF^ inhibition elicits consummatory behaviors directed at inanimate objects such as a wooden block and inhibition of mesencephalic trigeminal area (Me5) projections evokes rhythmic jaw movements. The activity of these neurons is decreased during food consumption and increases when food is in proximity but not consumed. Activity is also increased in obese animals and after leptin treatment. VMH^BDNF^ neurons receive monosynaptic inputs from both agouti-related peptide (AgRP) and proopiomelanocortin (POMC) neurons in the arcuate nucleus (Arc) and constitutive VMH^BDNF^ activation blocks the orexigenic effect of AgRP activation. These data delineate an Arc→VMH^BDNF^→Me5 circuit that senses the energy state of an animal and regulates consummatory behaviors in a state dependent manner.

## Introduction

Understanding the principles of how innate behaviors are generated has been a long-standing subject of intense theoretical and experimental inquiry but the precise neural mechanism underlying behavior selection are largely unknown (Lorenz, Tinbergen, Sherrington)^1–3^. Feeding is a typical innate behavior in which an array of sensory and interoceptive inputs are integrated to generate an adaptive behavioral response. However, because these many inputs typically change the probability of feeding, in contrast to a reflex, behavior initiation is context dependent^4^. While many brain areas and neuronal types have been identified that modulate feeding behavior, mapping a complete circuit linking relevant sensory and interoceptive inputs to consummatory behaviors is important for understanding how feeding is regulated. We thus sought to identify a neural pathway that directly connects interoceptive signaling to premotor centers that control consumption. Identifying the essential components of such a circuit would be informative in its own right and potentially provide a framework for investigating how top-down signals regulate innate behaviors^2^. Here we report the components of a simple circuit regulating feeding that directly connects leptin signaling in the arcuate nucleus (Arc) to the jaw premotor area of the mesencephalic trigeminal area (Me5) that control food consumption.

The elements of this circuit emerged from studies aimed at identifying which population(s) of BDNF neurons regulate food intake and body weight. A role for BDNF neurons in feeding has been shown in studies of mice and humans carrying mutations in brain-derived neurotrophic factor (BDNF) or its receptor tyrosine kinase B (TrkB)^5–11^. Heterozygous BDNF or TrkB mutations cause overconsumption and massive obesity in animals and humans. In addition, GWAS studies ^12,13^ have also suggested a role for BDNF in the development of obesity. However, because BDNF is widely expressed, the neural mechanism(s) responsible for the obesity and hyperphagia seen with defects in BDNF signalling are unknown. Because BDNF mutations cause obesity, we initially set out to identify which BDNF neurons normally restrict overfeeding in animals fed a highly palatable diet. We found that a discrete subpopulation of BDNF neurons in the ventromedial hypothalamus (VMH), but not elsewhere, are activated in diet induced obese (DIO) mice fed a high fat diet (HFD). We then set out to define the function of these neurons including the identification of their inputs and outputs. Here we report that VMH^BDNF^ neurons act as a key node regulating food consumption that directly links neurons in the arcuate nucleus that receive interoceptive inputs to premotor sites in the brainstem that regulate jaw movements and consummatory behavior.

## Results

### BDNF expressing neurons in the VMH regulate food intake and body weight

Mutations in BDNF or TrkB, its receptor, cause extreme obesity in mice and humans indicating that neurons that express these genes normally function to restrict food intake and weight gain. However, the identity of the BDNF neurons that regulate feeding and weight is not knows. In order to elucidate the neural mechanism, we reasoned that the key subpopulation(s) of brain-derived neurotrophic factor (BDNF) expressing neurons would be activated when animals are placed on a HFD and gain body weight. This was investigated by identifying those BDNF neurons that show increased c-fos expression, an activity marker, in response to a HFD in BDNF-cre mice^14^ mated to Rosa26^fsTRAP^ reporter mice expressing a cre dependent GFP. Brain sections from HFD fed and chow fed mice were co-stained for both GFP (BDNF) and c-fos. This revealed a 50-fold increase in the number of BDNF neurons that express c-fos in the central and dorsomedial regions of the VMH (Fig.1a,b). Indeed, there was a nearly complete overlap between GFP and cFos in VMH with 98% of the neurons from DIO mice expressing c-fos also expressing BDNF (Fig. 1c). Furthermore, in situ hybridization for Sf1 and BDNF revealed that approximately half of the BDNF neurons in the dorsomedial VMH do not express SF1 (Extended Data Fig. 1a-c), confirming previous RNAseq^15^ and immunohistology^16^studies showing that the BDNF expressing neurons are a distinct subpopulation of the neurons in this nucleus. The VMH^BDNF^ neurons show limited overlap with steroidogenic factor 1 (Sf1), a canonical marker for VMH which, as shown below, have different functional effects^17–19^. The data thus suggests that VMH^BDNF^ neurons represent a distinct subpopulation of VMH neurons that are specifically activated in animals fed a HFD. Further studies (Extended Data Fig.1d,e) revealed that 99% of the VMH^BDNF^ neurons colocalize with vesicular glutamate transporter 2 (Vglut2) and are thus glutamatergic. To directly test the possibility that these neurons normally restrict food intake and body weight, we analysed the effect of selectively ablating VMH^BDNF^ neurons in mice fed chow and high fat diets.

**Figure 1.**
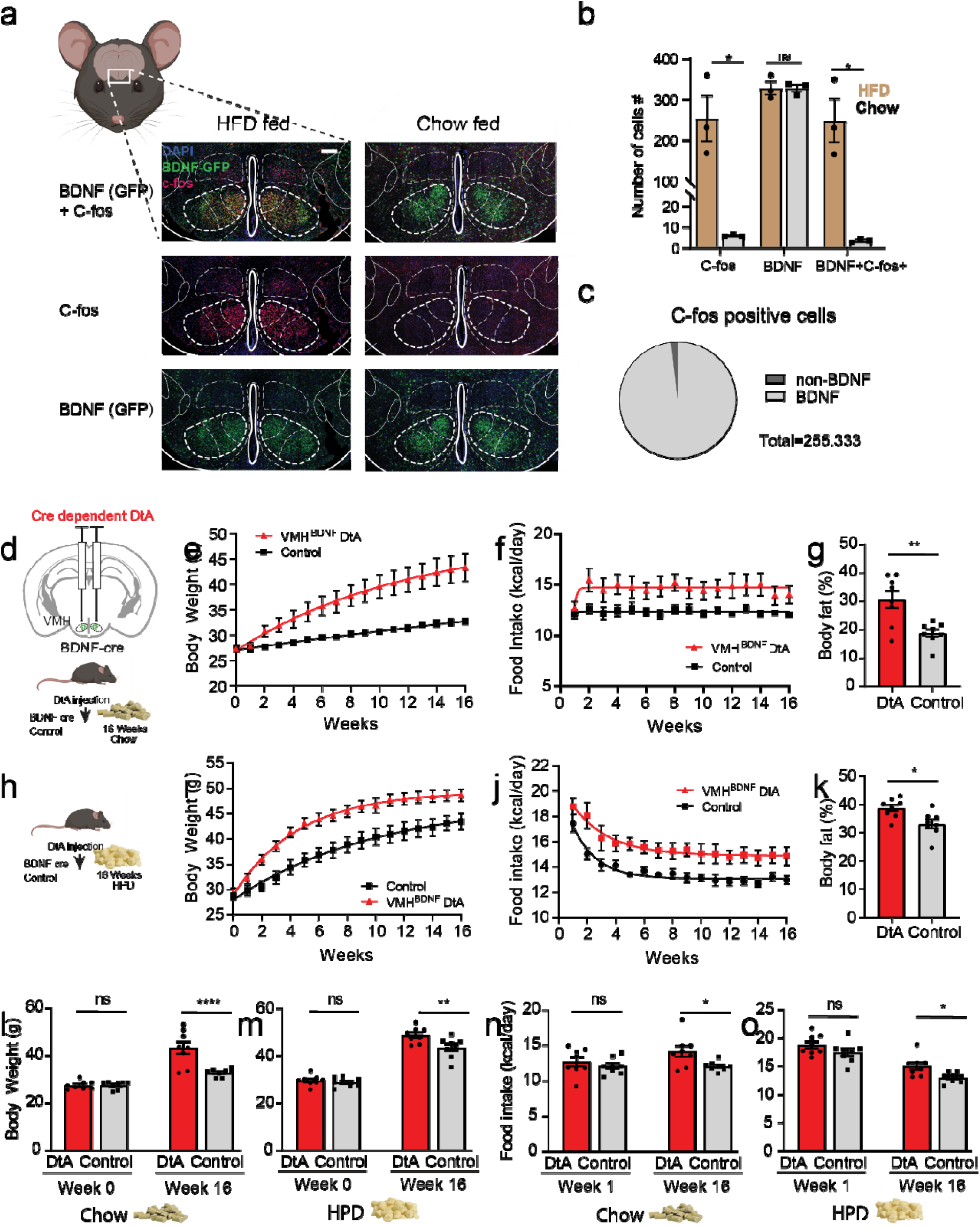
VMH^BDNF^ neurons suppress food intake and body weight. **a,** Schematic of a coronal brain slice containing the VMH and representative example of a hypothalamus section with immunolabelling for c-fos (red) and BDNF-GFP (green) after 16 weeks of HFD. Scale bar represents 200μm. **b,** Quantification of the number of c-fos positive and BDNF-GFP cells in the VMH of 3 mice. C-fos: 250.7±56.19 cells in HFD fed mice vs. 5.33±0.33 cells in sections from chow fed control mice, P=0.031868, BDNF: 325.333±16.58 cells in HFD fed mice vs. 325.33±9.207 cells in sections from chow fed, P= >0.999999, BDNF+cfos: 246.0±53.873 cells in HFD fed mice vs. 3.0±0.577 cells in sections from chow fed, P=0.031868, Multiple, paired t-tests with Holm Sidak method. **c,** Quantification of c-fos positive cell expressing BDNF-GFP. **d,** schematic of bilateral DtA injection and experimental timeline for mice fed chow ad lib. **e,** weekly time course of body weight and calorie intake **(f)** of chow fed DtA (n= 8 mice) and control mice (n= 8 mice). **g,** Comparison of % body fat between DtA and control. 30.59±3.07% for DtA vs 18.85±1.37% for control, P=0.006, Unpaired t test with Welch’s correction **h,** experimental timeline for mice fed HPD ad lib. **i,** weekly time course of body weight and calorie intake (**j**) of HPD fed DtA (n= 8 mice) and control mice (n= 8 mice). **k,** Comparison of % body fat between DtA and control on HPD. 38.66±1.22% for DtA vs 32.95±1.55% for control, P=0.0125, Unpaired t-test with Welch’s correction **l**, Body weight comparison between DtA and control mice before DtA ablation and 16 weeks post-surgery. For chow 43.28±2.79g for DtA vs 32.68±0.60g for controls, P <0.0001, Two-way RM ANOVA with Šídák’s multiple comparisons test. and **m**, HPD fed mice: 48.65±1.21g for DtA vs 43.42±1.57g for control, P =0.0055, Two-way RM ANOVA with Šídák’s multiple comparisons test**. n,** Comparison of average daily calorie intake between DtA and control mice 1-and 16-weeks post-surgery for chow: 12.04± 0.24kcal for DtA vs 14.14± 0.80kcal for control, P =0.0303, Two-way RM ANOVA with Šídák’s multiple comparisons test and **o**, HPD fed (right) mice. week 1: 18.77±0.65kcal in DtA vs 17.45±0.68kcal in controls, P=0.2355, Two-way RM ANOVA with Šídák’s multiple comparisons test week 16: 14.99± 0.65kcal for DtA vs 12.98± 0.32kcal for control, P =0.047, Two-way RM ANOVA with Šídák’s multiple comparisons test

VMH^BDNF^ neurons were ablated by injecting an adeno-associated virus (AAV) expressing a floxed diphtheria toxin A subunit (DtA) construct^20^ into the VMH of BDNF cre mice (Fig. 1d-o). Chow fed mice with DtA ablation (Fig.1d-g) ate significantly more than controls (Fig.1n) resulting in a significantly increased body weight (Fig.1l) and body fat content (Fig.1g,). We next ablated VMH^BDNF^ neurons in animals fed a highly palatable diet (HPD) (Fig. 1h-k). Initially, mice with an ablation of VMH^BDNF^ neurons showed overfeeding similar to that of control mice (Fig. 1j and m,). However, while food intake in control animals fed a HPD gradually decreased over time as weight increased, animals with VMH^BDNF^ neuron ablation continued to consume significantly more food for an extended period (Fig.1j and m). When fed a HFD, mice with a VMH^BDNF^ ablation ultimately stabilized at a significantly higher level than that of control mice (Fig.1i and o). Mice with VMH^BDNF^ ablation also showed a significantly greater level of adiposity (Fig.1k). These results show that VMH^BDNF^ neurons normally function to restrict overfeeding and obesity in animals fed a HPD or chow diet.

### VMH^BDNF^ neurons bidirectionally control food intake

To investigate whether VMH^BDNF^ neuron activity regulates food intake, we used optogenetics to either inhibit or activate these neurons. AAV strains encoding channelrhodopsin (ChR)^21^ (Fig.2a-d) or Guillardia theta anion channelrhodopsin (GtACR)^22^, (Fig.2e-h) were injected into the VMH of BDNF-cre mice and food intake was measured in response to light. To investigate whether activation of VMH^BDNF^ neurons suppresses feeding, we fasted mice overnight and then presented them with a chow pellet (Fig. 2c). Without light activation, control and ChR mice consumed similar amounts of food during the first 30 min. However, VMH^BDNF^ neuron stimulation at 2Hz completely inhibited food intake by 99.8 % which returned to control levels when light stimulation ceased. Thus, VMH^BDNF^ neuron activation can entirely suppress the hunger-induced, drive to eat. To test whether VMH^BDNF^ neuron activity is also sufficient to suppress hedonic feeding, we presented sated chow fed mice with a highly palatable pellet with high sugar and high fat content (Fig.2d). Control and ChR mice consumed similar amounts at baseline but optogenetic activation of VMH^BDNF^ neurons at 2Hz significantly decreased intake of the highly palatable pellets by 76 %. We also found that in the period after light activation ceased, the mice in which VMH^BDNF^ neurons were activated now consumed even more HPD than controls. These data show that VMH^BDNF^ neural activation significantly diminishes both homeostatic and hedonic feeding.

**Figure 2.**
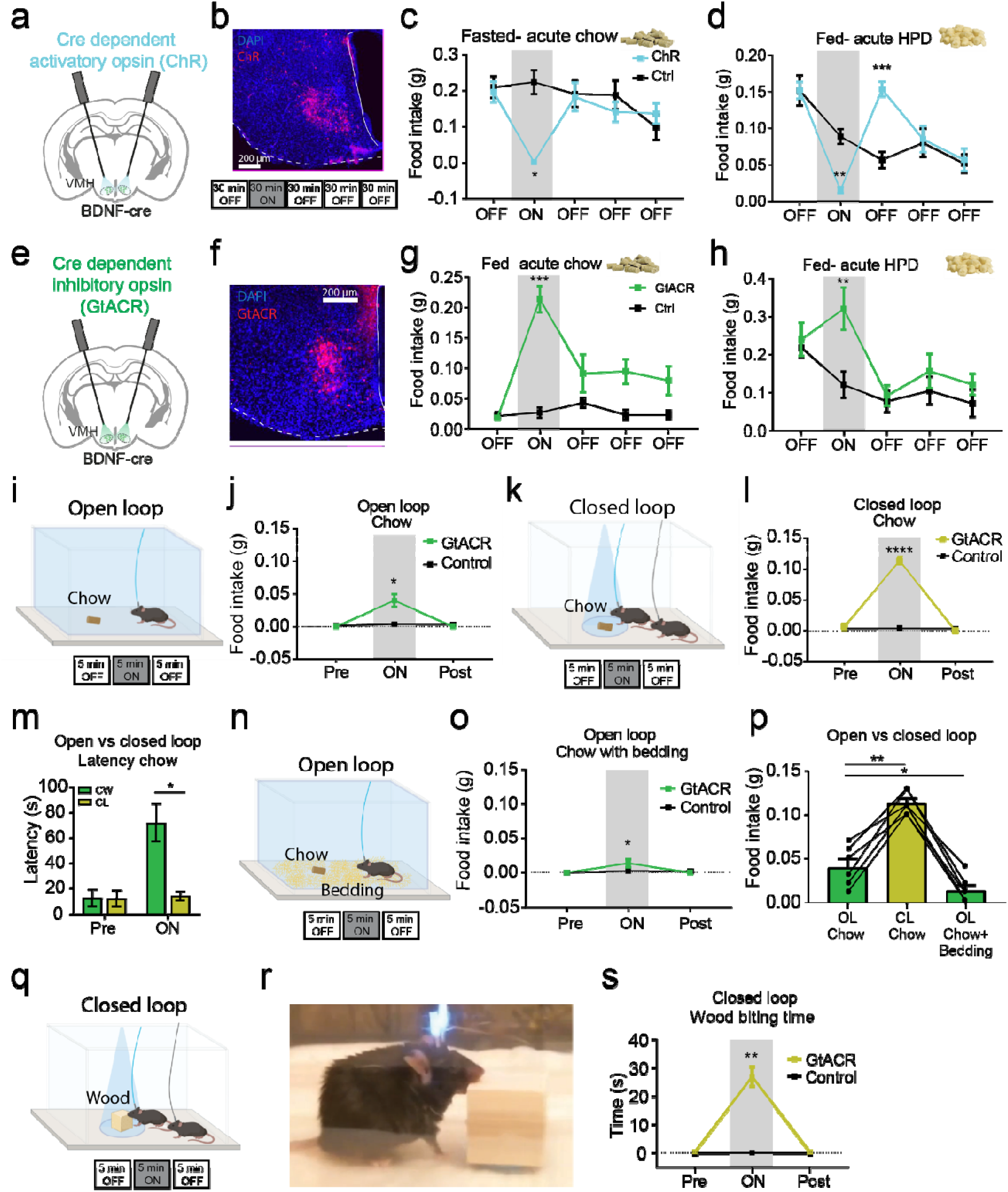
VMH^BDNF^ neuron activity bidirectionally controls food consumption. **a,** Schematic of ChR expression and optic fiber placement. **b,** Example of ChR expression with experimental timeline. **c**, Food intake of fasted ChR (n=6) and control mice (n=6) tested with acute chow (for ChR mice 0.002±0.002g vs 0.233±0.033g for controls, P=0.056, Two-way RM ANOVA with Šídák’s multiple comparisons test) and **d,** fed mice with acute HPD. (For ChR mice 0.015±0.004g vs 0.088±0.010g for controls, P=0.0029, Two-way RM ANOVA with Šídák’s multiple comparisons test) **e,** Schematic of GtACR expression and optic fiber placement. **f,** example of GtACR expression. **g**, Food intake of chow ad lib fed GtACR (n=6) and control mice (n=6) tested with acute chow and **h,** with acute HPD for ChR mice 0.153±0.010g vs 0.057±0.011g for controls, P= <0.0001, Two-way RM ANOVA with Šídák’s multiple comparisons test). **i**, Schematic of open loop optogenetic inhibition and experimental timeline. **j,** Quantification of food intake of GtACR (n=6) and control mice (n=6). **k,** Schematic of closed loop optogenetic inhibition and experimental timeline. **l,** Quantification of food intake of GtACR (n=6) and control mice (n=6). **m**, Comparison of the latency to approach the chow pellet in open and closed inhibition for GtACR mice. 72.00±14.83s in OL vs 14.14±3.49s in CL P= 0.0139, Two-way RM ANOVA with Šídák’s multiple comparisons test. **n,** Schematic of open loop optogenetic inhibition with bedding present and experimental timeline. **o,** Quantification of food intake of GtACR (n=6) and control mice (n=6). **p**, Comparison of food intake during optogenetic inhibition of 6 GtACR mice in the open loop, closed loop and open loop with bedding set up.,0.04±0.010g for OL vs 0.113±0.006g for CL, P=0.0012, 0.04±0.010g for OL vs 0.013±0.006g for OL with bedding, P=0.018, Mixed-effects model (REML) with Dunnett’s multiple comparison **q**, Schematic of closed loop optogenetic inhibition with a wood block and experimental timeline. **r**, Example of consummatory biting behavior displayed by mice upon VMH^BDNF^ neuron inhibition. **s,** Quantification of % of time spent biting the wood block for GtACR (n=6) and control mice (n=6). 26.84±3.423s for GtACR mice vs 0.001±0.001s for controls, P=0.00166, Two-way RM ANOVA with Šídák’s multiple comparisons test

We next tested the effect of VMH^BDNF^ neuron inhibition by measuring food intake in fed BDNF-cre animals expressing GtACR in VMH neurons. While the baseline intake of the control and GtACR mice was the same, optogenetic inhibition in the GtACR mice significantly increased food intake by 1183 % during the 30-minute testing period (Fig. 2g). To test whether inhibiting VMH^BDNF^ neuron activity would also increase the intake of HPD, we measured food intake after presenting mice with a HPD pellet. Initial intake prior to optogenetic inhibition was similar between control and GtACR mice and light induced neural inhibition again increased consumption to 133% (Fig.2h).

We next evaluated our hypothesis that VMH^BDNF^ neurons are activated by obesity to suppress further weight gain (see data in Fig. 1a-c), by measuring food intake during VMH^BDNF^ inhibition in DIO mice that had been fed a HPD for 4 weeks. Consistent with the studies of VMH^BDNF^ neuron inhibition in lean mice, optogenetic inhibition of these neurons using GtACR in the obese mice increased consumption of both chow (706 %) and HPD (1279 %) pellets during the epochs of light exposure (Extended Data Fig.2 a-d).

In these studies, we tested the effect of VMH^BDNF^ inhibition in animals before and after they were fed HPD. We noted that at baseline, prior to optogenetic inhibition, when the DIO mice were later presented with chow, they consumed smaller amounts then they had before they fed the HPD (Extended Data Fig 2e). One potential explanation for this finding is that the DIO animals fed HPD devalue chow ^23,24^. However, we found that at baseline before VMH^BDNF^ inhibition the DIO animals also consumed less of the HPD (Extended Data Fig 2f). We thus hypothesized that this reduced consumption of the HPD and chow after obesity has developed might be caused by hyperactive VMH^BDNF^ neurons (as suggested by the increased cFos expression). If true, optogenetic inhibition of VMH^BDNF^ neurons should have a greater quantitative effect in DIO vs lean animals. Consistent with this, we found that optogenetic inhibition of VMH^BDNF^ neurons using GtACR resulted in a greater quantitative increase in food intake in DIO compared to lean animals (Extended Data Fig. 2g). We interpret this to mean that VMH^BDNF^ neurons become more active once animals develop Diet Induced Obesity at which point, they then restrict overeating. (We also imaged these neurons and indeed find increased Ca^2+^ signal in VMH^BDNF^ neurons from DIO mice, see below). These studies show that VMH^BDNF^ neurons normally act to restrict intake of both chow and palatable food by DIO animals.

We then assessed whether BDNF itself contributes to these effects by inhibiting BDNF signaling in DIO mice using a TrkB^F616A^ knockin mouse carrying a point mutation that renders the receptor sensitive to an allele specific kinase inhibitor (1-NM PP)^25^. Homozygous TrkB^F616A^ mice were fed a high fat diet (HFD) after which they were treated with 1-NM PP1. Mice treated with 1-NM PP1 showed a significant increase of body weight of 17 % (Extended Data Fig. 2h) and food intake of 27% (Extended Data Fig. 2i, j). These data show that BDNF signalling is also required to suppress food intake and weight gain in HPD fed mice after obesity has developed.

To further explore the mechanism by which VMH^BDNF^ neurons can control feeding, we tested whether VMH^BDNF^ neuron activity alters valence which would suggest a possible role in motivation. However, in an operant conditioning task, VMH^BDNF^ neuron activation failed to entrain a preference for self-inhibition (Extended Data Fig.2 h-j) and also failed to entrain a preference in a flavour conditioning assay (Extended Data Fig.2 k-m). These studies suggest that the effect of VMH^BDNF^ neurons on feeding is not mediated by a general effect on valence. An alternative possibility is that VMH^BDNF^ neurons regulate the consummatory rather than the appetitive phase of feeding. We tested this by analysing whether VMH^BDNF^ neuron inhibition influenced food approach vs. food consumption after optogenetic inhibition of VMH^BDNF^ neurons using two different paradigms: i. Constant photoinhibition for 5 min in an open loop (OL) system (Fig.2i,j) ii. inhibition in a closed loop (CL) system in which light was delivered only when the head of the mouse was within 3 cm of a chow pellet (Fig.2k, l). We found that the CL configuration led to a ∼3 times greater food intake than did OL inhibition (Fig.2l). Consistent with this, the latency to approach the pellet was also increased 5-fold in the OL paradigm showing that VMH^BDNF^ neuron inhibition decreases the rate at which mice approach food (Fig.2m). We also noted that OL VMH^BDNF^ neuron photoinhibition often triggered consummatory behaviour targeted at objects in immediate proximity to the mouse including the wall and bare floor of the cage. Thus, VMH^BDNF^ neuron photoinhibition not only reduced food approach but appeared to trigger motor behaviours associated with consumption. To characterise this further we studied the effect of VMH^BDNF^ neuron photoinhibition in the OL system in the presence of corn cob bedding (Fig.2n, o). We observed that animals would often chew the bedding during VMH^BDNF^ neuron photoinhibition and that the presence of bedding further decreased food intake compared to OL stimulation in a bare cage (Fig.2p). These data show that VMH^BDNF^ neuron inhibition drives feeding most effectively when food is in proximity to the animal and suggests the possibility that VMH^BDNF^ neurons regulate motor programs required for consumption independent of caloric value. To further evaluate this, we repeated the CL study, but now provided the mice with a wooden block instead of a chow pellet (Fig. 2q-s). VMH^BDNF^ neuron inhibition in the CL led mice to spend ∼9% of the trial duration engaged in biting the wooden block (Supplementary Video S1,). Taken together, these results suggest that inhibition of VMH^BDNF^ neuron activity increases food intake by triggering consumption of whatever objects are in proximity (such as a wood block or corn cob bedding) rather than by altering valence or food seeking. This finding is analogous to that seen after activation of central amygdala projections to the reticular formation, the motor pattern generator for killing bites^26^. Moreover, it raises the possibility that VMH^BDNF^ neurons represent a distal node, downstream of hedonic and homeostatic drives, in the neural circuit regulating feeding that controls the motor components of consumption such as biting and chewing. To test whether VMH^BDNF^ neuron inhibition would also elicit other types of biting such as aggressive biting, we placed mice expressing GtACR in VMH^BDNF^ neurons in an open field box with a female mouse (Extended Data Fig.2n, o) but failed to observe any biting attacks or aggression during photoinhibition. We also noticed that inhibition dramatically reduced the amount of time mice spent interacting with the females 86%. In this study, we again observed the same motor behaviors (i.e; biting) that in this case were directed at the wall and floor of the cage as was previously observed after open loop inhibition (Fig. 2j).

### VMH^BDNF^ neural activity is inversely correlated with feeding and tuned to energy state

The potent effect of VMH^BDNF^ neurons to reduce feeding suggested that the activity of these neurons might be decreased as they consumed food and thus permissive for feeding. We measured the in vivo activity of VMH^BDNF^ neurons using fiber photometery to monitor intracellular calcium levels when animals consumed or rejected food. BDNF-cre mice received injections of a cre dependent AAV encoding GCamp6s into the VMH (Fig. 3a) and calcium signals were measured during bouts of feeding. Time locked recordings showed a significant 4 % decrease in VMH^BDNF^ neuron activity coinciding with consumption of a chow pellet (Fig.3b, c, Supplementary Video S2,). This is consistent with the optogenetic results and suggests that VMH^BDNF^ neuron activity gates feeding behaviour and needs to be inhibited for consumption to commence. To test whether VMH^BDNF^ neuron activity is influenced by food palatability, we recorded their response during consumption of sucrose treat pellets which revealed a decrease of similar magnitude as was seen after chow consumption (Fig. 3b,c).

**Figure 3.**
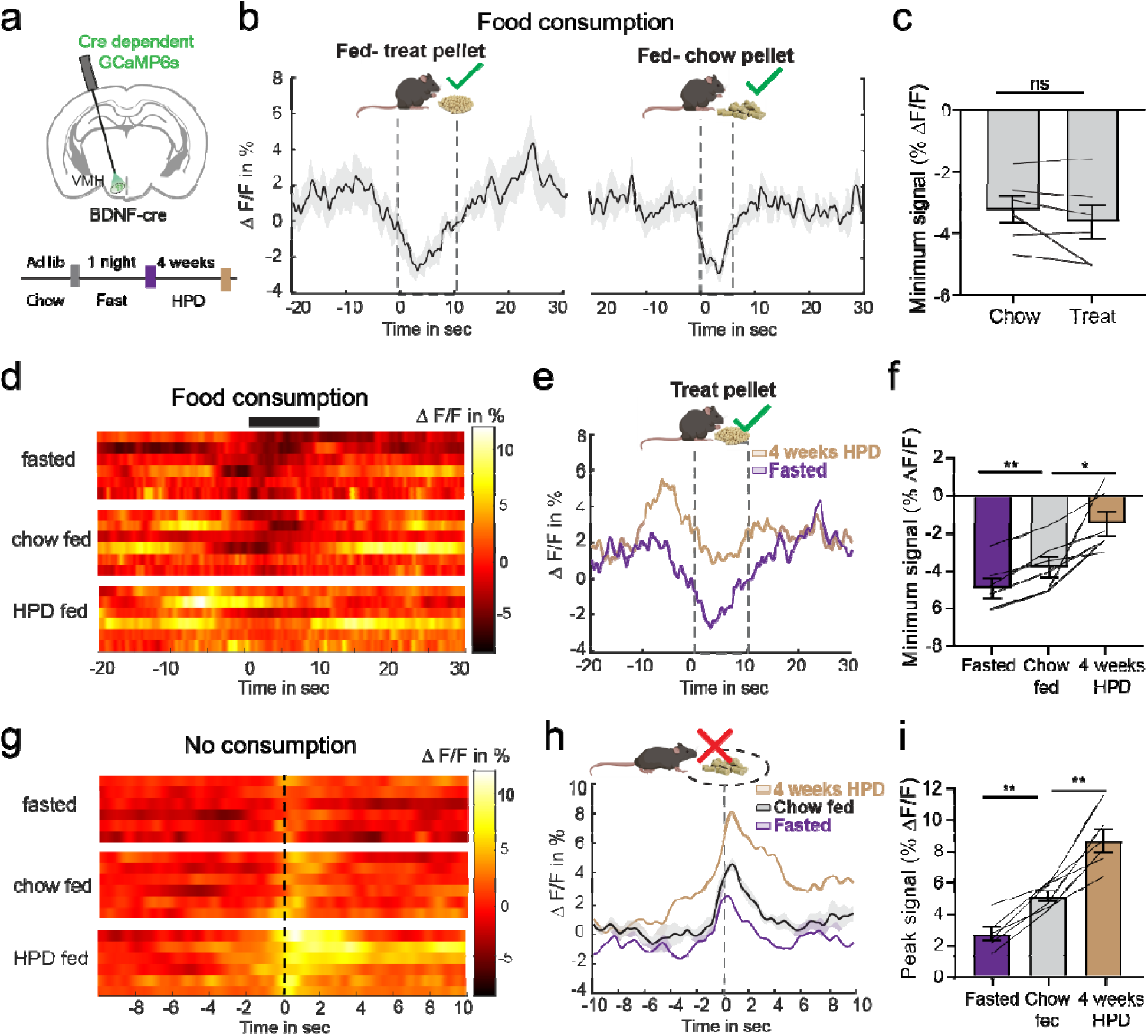
VMH^BDNF^ neuron activity is decreased during food consumption and tuned to energy state. **a**, schematic of a coronal brain slice with GCaMP expression and fiber placement and below recording timeline for energy state manipulations. **b**, average photometry trace ± sem (n=6 mice) aligned to a consumption bout of, left, a treat pellet in chow ad lib fed mice and right, a chow pellet. **c**, comparison of the minimum signal between chow and treat pellet consumption (n=6 mice). −3.674±0.5355% for sucrose treat vs −3.226±0.429% for chow, P=0.1501, Paired t-test. **d**, heatmap of average photometry recordings per mouse (n=6 mice) aligned to a bout of treat pellet consumption. **e**, average overall photometry trace ± sem (n=6 mice) aligned to a bout of treat pellet consumption in fasted (purple) and 4 weeks HPD fed (beige) mice. **f**, comparison of the minimum signal during treat consumption. −3.674±0.5355% in lean vs −4.792±0.5139 % in fasted (P=0.0013), lean vs −1.396±0.640% in DIO (P=0.0356), Mixed-effects model (REML) with Holm-Šídák’s multiple comparisons test. **g**, heatmap of average photometry recordings per mouse (n=6 mice) aligned to food approach without consumption. **h,** overall average overall photometry trace ± sem (n=6 mice) aligned food approach without consumption in fasted (purple) and 4 weeks HPD fed (beige) mice. **i**, comparison of the maximum signal. 5.149±0.3302% in lean vs 2.719±0.4372 % in fasted (P=0.0020), lean vs 8.703±0.7414% in DIO (P=0.0036), Mixed-effects model (REML) with Holm-Šídák’s multiple comparisons test.

Having established that VMH^BDNF^ neuron activity is reduced during the consumption of chow and sucrose pellets, we investigated the effects of alterations of energy balance. Neural activity was analysed in three sets of conditions: after an overnight fast and in mice fed a chow or a HPD for 4 weeks (Fig 3a. Baseline activity of these neurons in an empty home-cage showed no difference among these three conditions (Extended Data Fig. 3 a,b). However, recordings of the activity of VMH^BDNF^ neurons during bouts of feeding revealed a ∼60% higher level of activity of VMH^BDNF^ neurons in the DIO mice vs. chow fed animals (Fig. 3d-f). In addition, the activity in VMH^BDNF^ neurons was ∼30% lower in fasted vs. chow fed animals. We also noticed that when chow fed animals approached a chow pellet within a radius of 2 cm but didn’t consume it, the calcium signal in VMH^BDNF^ neurons significantly increased (Fig. 3g-i, Supplementary Video S3). There was also an increase in VMH^BDNF^ activity when fasted mice approached but did not consume the food. However, in this case the increased activity was only half of what was observed when fed mice approached but did not consume the pellet and only 30% of what was seen when DIO mice did not consume it (Fig. 3i). In aggregate, these data show that VMH^BDNF^ neural activity is inversely correlated with food consumption and that activity responses are tuned to the animal’s energy state.

### VMH^BDNF^ neurons are downstream of Arc^POMC^ and Arc^AgRP^ neurons

Our finding that VMH^BDNF^ neural dynamics is altered by energy state, i.e. in fed, fasted and DIO, raised the possibility that they might respond to the adipocyte derived hormone leptin^27,28^. This was tested using fiber photometry recordings after leptin vs. saline injections into fasted animals (Fig.4a-d). Leptin increased VMH^BDNF^ neural activity by ∼80% when animals approached but did not consume a chow pellet (Fig 4d) and also increased it by ∼ 25% when animals consumed a treat pellet in comparison to saline injected mice (Extended Data Fig 4a-c,). To confirm that these neurons contribute to leptin’s effect on food intake we treated animals with leptin, ip 3mg/kg, after VMH^BDNF^ neural ablation with DtA (Extended Data Fig.3 d-f). The effect of leptin to reduce food intake was significantly reduced in animals in which VMH^BDNF^ neurons were ablated. To test whether VMH^BDNF^ neurons can also respond to signals besides leptin, we crossed BDNF-cre mice to leptin deficient ob/ob mice and injected DtA into the VMH (Extended Data Fig.3 g-i). ob/ob mice with VMH^BDNF^ ablation showed significantly increased food intake and became more obese. Because ob/ob mice lack leptin, these data indicate that VMH^BDNF^ neurons also sense additional satiety signals besides leptin.

**Figure 4.**
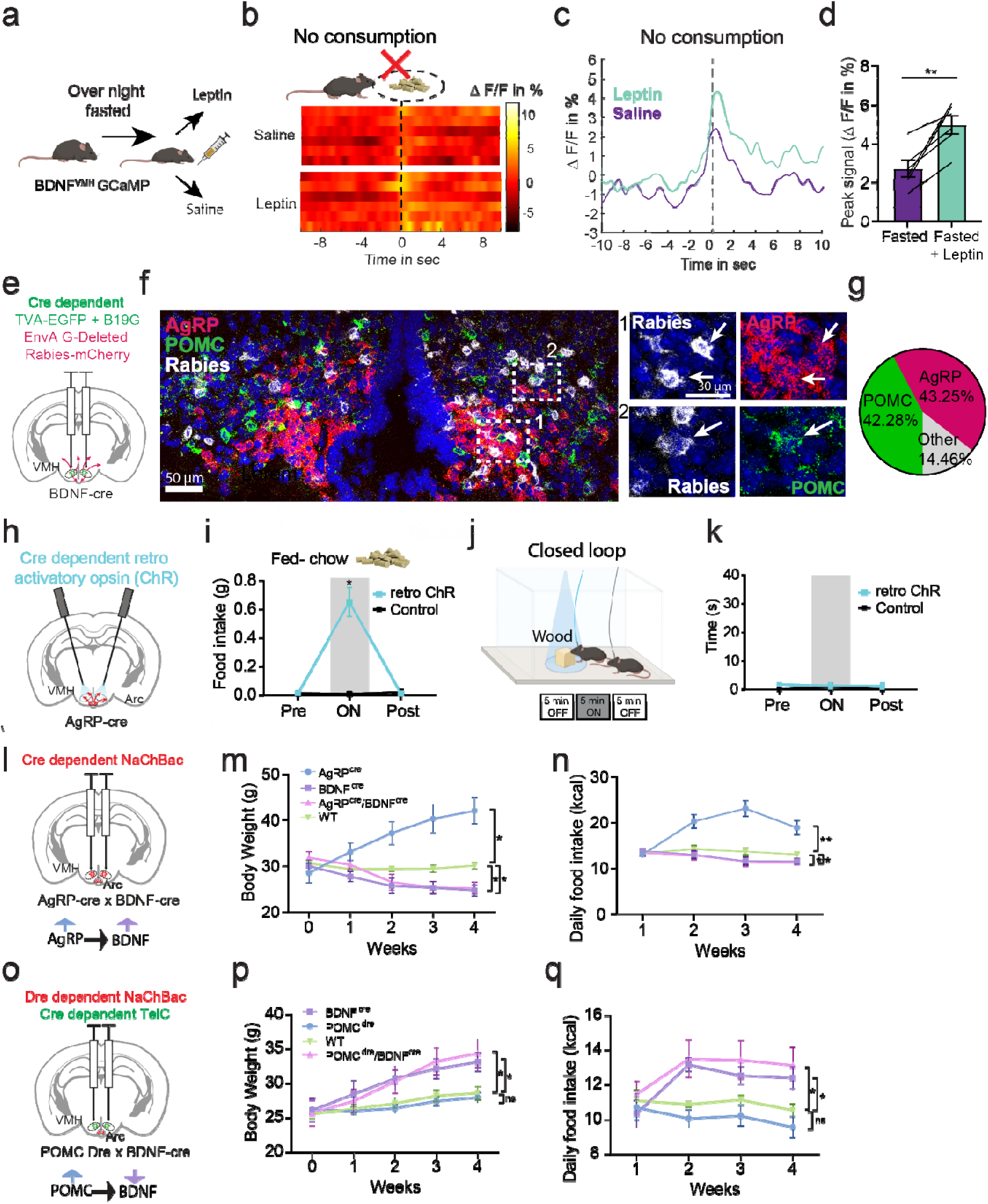
VMH^BDNF^ neurons are anatomically and functionally downstream of AgRP and POMC neurons. **a,** schematic of experimental design. **b,** heatmap of average photometry recordings per mouse and **c,** overall average photometry trace (±sem) (n=6 mice) aligned to food approach without consumption of mice injected with saline (purple) or leptin (green). **d,** comparison of the peak photometry signal: 2.719±0.4372% saline injected vs in 5.013±0.4727% leptin injected, P=0.0054, Paired t-test. **e,** Left, schematic of G-deleted pseudotyped rabies and helper AAV expression. **f**, representative example (n=3 mice) of a coronal Arc section with in situ hybridization for Rabies (white), AgRP (red) and POMC (green) and high magnification examples of the boxed area in (1) and (2). **g**, Quantification of average overlap of rabies labelled cells with AgRP and POMC in the Arc (n=3 mice). **h**, schematic of retrograde ChR expression in Arc and optic fiber placement in VMH. **i**, food intake of ad lib fed ChR (n=4) and control mice (n=4) tested with acute chow. **j**, schematic of closed loop optogenetic inhibition with a wood block and experimental timeline. **k**, quantification of % of time spent biting the wood block for ChR (n=4) and control mice (n=4). **l**, schematic of bilateral expression of cre dependent NaChBac in Arc and change in neural activity. **m**, weekly time course of body weight: AgRP cre 42.06 ±2.85g, BDNF cre 24.77±1.27g, AgRP cre/BDNF cre 25.21±1.32g, control mice 30.13±1.82g and calorie intake (**n**) of AgRP cre (n= 6 mice)16.86 ±2.06 kcal/day,BDNF cre (n=6mice) 11.53±0.76kcal/day, AgRP/BDNF cre (n=6mice) 11.37±0.55kcal and control mice (n= 7 mice) 13.14±1.29kcal/day after NaChBac injection. **o,** schematic of bilateral expression of dre dependent NaChBac in Arc and cre dependent TelC in VMH and respective change in neuronal activity. **p**, weekly time course of body weight:BDNF cre 33.75±1.40g, POMC dre 28.02±0.69g, POMC dre/BDNF cre 35.48±1.85g, control mice 28.78±2.40g and (**q**) calorie intake of POMC dre (n= 6 mice) 9.585±0.60kcal/day, BDNF cre (n=7mice) 12.42±0.48kcal/day, POMC dre/BDNF cre (n=7 mice) 13.22±0.80kcal/day and control mice (n= 7 mice) 10.56±0.98kcal/day after AAV injection.

Prior anatomic^29^ and single cell RNA sequencing studies^15^ have indicated that VMH^BDNF^ neurons do not express leptin receptors, suggesting that leptin’s effect on these neurons is indirect. Therefore, we next determined whether VMH^BDNF^ neurons receive inputs from the arcuate nucleus (Arc) (Fig. 4e), a key site of action for multiple interoceptive signals including leptin. The inputs to VMH^BDNF^ neurons were labelled by monosynaptic tracing with pseudotyped rabies^30,31^. We injected helper AAVs and G-deleted Rabies-mCherry viruses into the VMH of BDNF-cre mice and observed prominent inputs from Arc where leptin responsive POMC and AgRP neurons are located (Fig. 4f,g). We next performed multichannel FISH using probes for rabies, POMC and AgRP which revealed that a large proportion of Arc inputs are from either POMC (42.28±7.95%) or AgRP (43.25±3.53%) neurons. In addition, FISH with a LepR probe revealed that many of the rabies labelled AgRP and POMC neurons also express the leptin receptor (Extended Data Fig.4j-m). Consistent with prior studies^32–34^, in situ hybridization also revealed that 93±2.29% VMH^BDNF^ neurons co-expressed both neuropeptide Y receptor 5 (NPY5R) and melanocortin receptor 4 (MC4R), suggesting that VMH^BDNF^ neurons are a point of convergence for Arc^POMC^ and Arc^AgRP^ outputs (Extended Data Fig. 5a,b).

We next investigated whether the AgRP projections to the VMH can drive feeding similar to the effect somatic activation of AgRP neurons in the Arc^35^. A cre-dependent retrograde AAV encoding ChR was injected into the VMH of AgRP cre mice and optogenetic fibers were implanted above the VMH (Fig.4h). AgRP neurons are GABAergic and optogenetic activation of the AgRP projections to the VMH increased food intake in chow fed mice (Fig.4i) but did not cause a decrease in food approach (Extended Data Fig. 5c-e). Moreover, closed loop activation targeted at a wood block did not elicit motor sequences of consummatory behavior (Fig.4j,k), suggesting that while projection specific activation regulates food intake, activation of these inhibitory inputs by themselves is not of sufficient intensity to activate motor programs of food consumption. This contrasts with the effect seen when inhibiting the soma of VMH^BDNF^ neurons. AgRP projections to the paraventricular hypothalamus (PVH) have also been reported to increase food intake via MC4R expressing neurons^36–38^ and retrograde tracing from VMH^BDNF^ neurons also showed inputs from the PVH. In addition, in situ hybridization of PVH neurons for MC4R after retrograde tracing revealed that VMH^BDNF^ neurons receive also monosynaptic inputs from PVH^MC4R^ neurons (Extended Data Fig.5f,g). Thus, melanocortin pathways neurons from multiple sites converge on VMH^BDNF^ neurons.

To assess whether VMH^BDNF^ neurons are functionally downstream of interoceptive neurons in the Arc, we simultaneously activated both Arc^AgRP^ and VMH^BDNF^ neurons by expressing NaChBac^39^ in one or both populations (Fig.4 l-n). NaChBac is a bacterial sodium channel that leads to constitutive neural activation. AgRP cre mice were crossed to BDNF cre mice and an AAV expressing a floxed NaChBac was stereotactically injected into the Arc and VMH (Fig.4l). Consistent with prior results^40^, activation of Arc^AgRP^ neurons alone led to marked hyperphagia and obesity. Constitutive activation of VMH^BDNF^ neurons alone significantly reduced food intake and body weight. However, simultaneous activation of both populations in double positive AgRP cre/BDNF cre mice fully recapitulated the effect seen with BDNF activation alone with these animals showing significantly decreased food intake and weight. These data show that VMH^BDNF^ neurons are functionally downstream of Arc^AgRP^ neurons and thus an important output site.

The functional relationship between Arc^POMC^ and VMH^BDNF^ neurons was tested by crossing POMC dre mice to BDNF cre mice and injecting a dre dependent NaChBac into the Arc and a cre dependent TelC virus into the VMH (Fig.4o). TelC prevents vesicular release and thus silences neurons. Consistent with effect of DtA ablation of VMH^BDNF^ neurons, BDNF cre mice expressing TelC alone showed significantly increased food intake and body weight (Fig.4p, q). Constitutive activation of Arc^POMC^ neurons led to a slight decrease of food intake and body weight which is consistent with prior studies^41^. Here again the phenotype of the POMC dre/BDNF cre double positive mice recapitulated the phenotype of mice with constitutive inhibition of VMH^BDNF^ neurons alone. These data show that VMH^BDNF^ neurons are also functionally downstream of Arc^POMC^ neurons. Overall, this set of experiments shows that VMH^BDNF^ neurons receive inputs from interoceptive neurons and are functionally downstream of them. We next determined where they project to and further assessed their effects consummatory behaviors.

### Projections from VMH^BDNF^ neurons to premotor areas of the jaw control food consumption

We mapped the projection sites of VMH^BDNF^ neurons by injecting an AAV with a Cre dependent mGFP-synaptophysin-Ruby gene into the VMH of BDNF-cre mice (Fig.5a-c). Fluorescence imaging of neurons expressing mGFP revealed dense projections to several known premotor sites in the brainstem that have previously been shown to send projections onto motor neurons of the jaw muscle and tongue including the mesencephalic nucleus (Me5), lateral paragigantocellular (LPGi) and the gigantocellular reticular nucleus alpha part (GiA) with only weak projections to the parvocellular reticular formation (PCRT) ^26,42–45^. We next assessed the function of each of these VMH^BDNF^ neuron→ premotor projections by optogenetically activating VMH^BDNF^ nerve terminals there. Optical fibers were implanted above the Me5, LPGi and GiA and PCRT in animals expressing ChR in VMH^BDNF^ neurons (Fig.5d-g) and feeding of a chow pellet was tested after an overnight fast. Projections to the LPGi and GiA were of interest since the LPGi-motor connection develops during weaning and might therefore be implicated in solid food consumption. However, activation had only a modest effect to reduce feeding by 37% whilst projections to the PCRT, which has been reported to stimulate the killing bite on crickets, failed to suppress feeding at all. In contrast, projections to Me5 significantly reduced feeding by 80 % which is similar to the effect we observed when stimulating VMH^BDNF^ soma. In addition, activation of Me5 projections also suppressed consumption of HPD feeding (Fig. 5h) and was specific for solid foods as it did not affect liquid diet licking (Extended Data Fiig, 6 c,d). Moreover, Me5 projections specifically control food consumption and not approach since inhibition of the terminals did not shorten the latency to approach a chow pellet (Extended Data Fig.6a,b).

**Figure 5.**
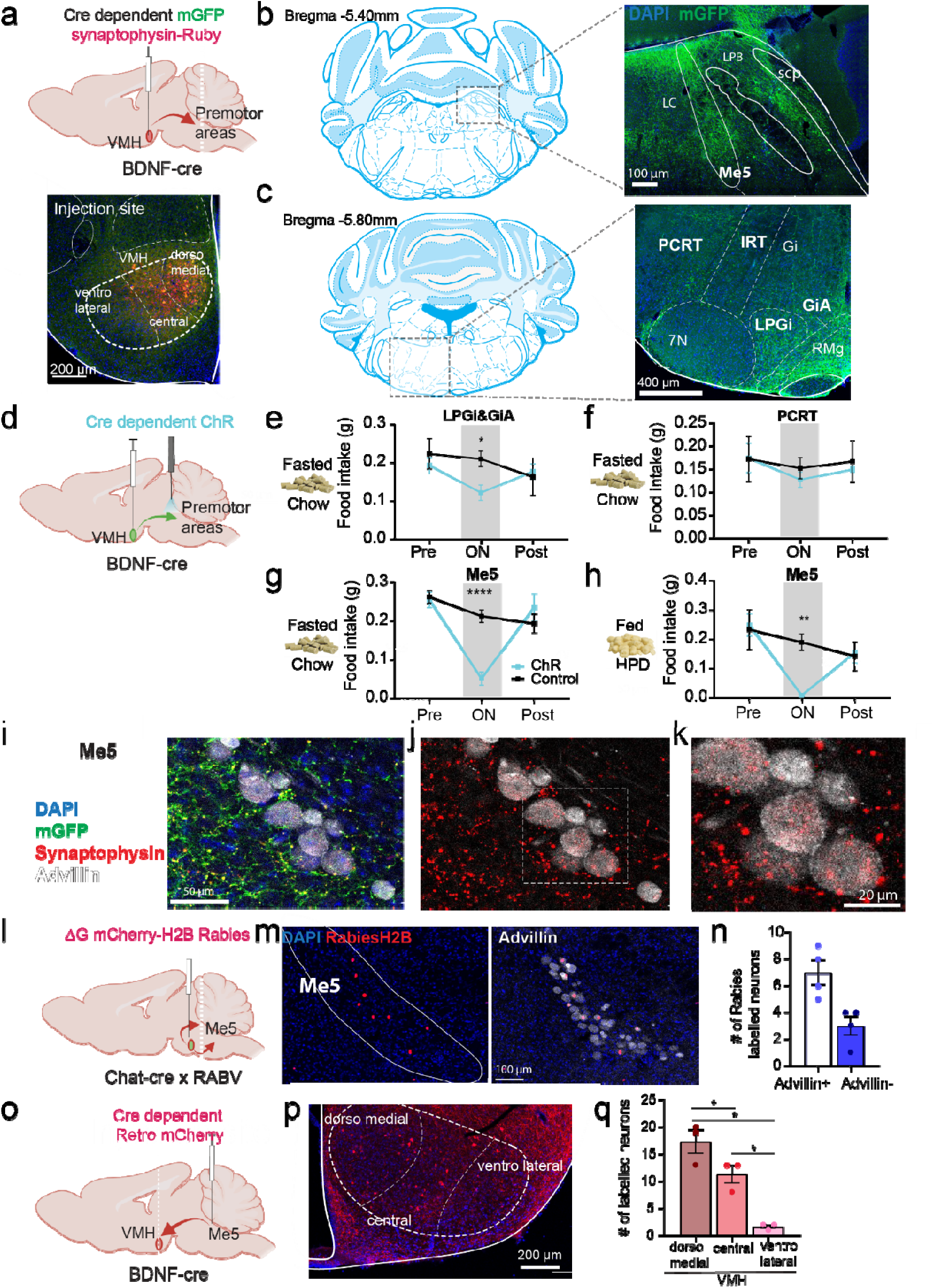
Activation of VMH^BDNF^ neuron projections to premotor areas. **a,** schematic of a sagittal brain with mGFP-synaptophysin-Ruby expression in VMH and level of sectioning. Below, representative image of injection site into VMH. **b,** Left, coronal section from brain atlas with dotted square indicating the imaged section on the right: representative (n=3 mice) coronal section of mGFP expression and DAPI (blue) of Me5.**c,** Left, coronal section from brain atlas with dotted square indicating the imaged section on the right: representative (n=3 mice) coronal section of mGFP expression and DAPI (blue) of PCRT and LPGi/GiA. **d**, schematic of a sagittal brain with ChR expression in the VMH and fiber placement above premotor areas. **e,** food intake of overnight fasted mice (n=5) with implants above LPGi&GiA tested with acute chow (ChR: 0.210±0.020g, Control: 0.122±0.020g, P=0.0444, Two-way RM ANOVA with Šídák’s multiple comparisons test). **f,** food intake of overnight fasted mice (n=4) with implants above PCRT tested with acute chow (ChR: 0.128±0.017g, Control: 0.153±0.023g, P=0.7976, Two-way RM ANOVA with Šídák’s multiple comparisons test). **g,** food intake of overnight fasted mice tested with acute chow (ChR: 0.05143±0.017g, Control: 0.2129±0.016g, P<0.0001, Two-way RM ANOVA with Šídák’s multiple comparisons test) and **(h)** ad lib fed tested with HPD ChR (n=7) and control mice (n=7) (ChR: 0.008571±0.003g, Control: 0.1900±0.028g, P= 0.0019, Two-way RM ANOVA with Šídák’s multiple comparisons test). **i,** Representative coronal image of Me5 after mGFP-synaptophysin-Ruby injection in VMH (see a) with projections in green, synaptophysin in red and immunofluorescent staining for advillin in white. **j**, same image as in (i) but with advillin (white) and synaptophysin (red) only. **k**, enlargement of the dotted square in j. **l**, schematic of a sagittal brain with ΔG mCherry-H2B Rabies injection into Mo5 and level of sectioning at Me5. **m**, Representative coronal image of Me5 with Rabies mCherry labelled nuclei in red and overlay with advillin immunofluorescent staining in white (right). **n**, quantification of rabies labelled neurons and their co-labelling with advillin: avillin+ 7±0.9129 neurons, avillin-3±0.7071 neurons. **o,** schematic of a sagittal brain with a cre dependent retro AAV mCherry injection into Me5 and level of sectioning at VMH. **p**, representative image of VMH with MCherry labelled BDNF neurons in red and DAPI. **q,** Quantification of mCherry labelled neurons in the dorsomedial (17.33±2.186 neurons), central (11.33±1.667 neurons) and ventrolateral (1.667±0.3333 neurons) part of VMH. dorsomedial vs central: P= 0.0271, dorsomedial vs ventrolateral: P=0.0425, central vs ventrolateral: P= 0.0425, Mixed-effects analysis with Holm-Šídák’s multiple comparisons test.

We next characterized the inputs and outputs from Me5 more fully. Me5 is a small nucleus and to further confirm that VMH^BDNF^ neuron project there, we performed immunofluorescence for advillin, a well-established marker for Me5 neurons^46^, together with Ruby-tagged synaptophysin introduced into VMH^BDNF^ neurons as above (Fig. 5 i-k). Dense synapses were found in Me5 and on advillin expressing cells as well as cells in Me5 that don’t express advillin. Monosynaptic retrograde rabies tracing from Chat-cre expressing neurons in thetrigeminal motor nucleus (Mo5) confirmed that in Me5 both advillin and non-advillin neurons project to premotor neurons in Mo5 (Fig. 5 l-n).

Next, we performed retrograde tracing from Me5 by injecting a retro AAV expressing a cre-dependent mCherry into the Me5 of BDNF cre mice. In line with the c-fos data (Fig.1a), we found that the majority of BDNF neurons projecting to Me5 are located in the ventromedial and central part of the VMH project (Fig 5p-r). We also investigated whether the other non-BDNF VMH populations project to Me5 using a ‘cre off’ strategy (Extended Data Fig.6 g-i). In this experiment, we used an AAV in which cre turns off a eYFP cassette such that it will be constitutively expressed in neurons that don’t express cre. A ‘cre-off’ AAV encoding eYFP-ChR was injected into the VMH of BDNF cre mice and the projection sites were ascertained. The non-BDNF neurons in VMH showed only minimal projections in Barrington nucleus and Locus Coeruleus and failed to show projections to Me5.

To investigate potential inputs to Me5 from extrahypothalamic areas with known premotor effects on jaw movements, we tested for projections from Central Amygdala (CeA). CeA neurons control the killing bite of crickets during prey hunting and have been reported to project to the PCRT ^26^. However, we failed to see any additional inputs to Me5 from Central Amygdala (using two anterograde tracing AAV1 viruses encoding either flp or cre ^47^. In this study, the flp expressing AAV1was injected into the CeA and the cre AAV1 was injected into the VMH of mice carrying both a cre dependent GFP and flp dependent tdTomato reporter (Extended Data Fig.6 j-l). In this study, CeA projections appear red and VMH projection neurons appear green. As before, the VMH projection neurons were localized primarily in Me5 while, consistent with previous report the CeA projections were seen in PCRT neurons^26^ and medial and lateral parabrachial nucleus next to Me5 in (Extended Data Fig. 6i,j). Our finding that the targets of the VMH^BDNF^ neurons are different from those from those from CeA is consistent with de Araujo’s finding that the CeA-> PCRT circuit does not alter food consumption.

### VMH^BDNF^ projections to Me5 regulate feeding and rhythmic jaw movements

We next investigated whether inhibition of the VMH^BDNF^ projections to Me5 recapitulated the effects seen during inhibition of VMH^BDNF^ soma (see Fig. 2 e-h). A Cre dependent AAV encoding eOPN3, an inhibitory mosquito-derived rhodopsin^22^, was injected into the VMH of BDNF-cre mice and implants were placed above the Me5 projection sites (Fig.6 a,b). Inhibition of the VMH^BDNF^ projections to Me5 increased food intake of both chow (863 %) and HPD diets (170 %) with a similar magnitude as was seen after inhibition of the soma (Fig.6 c,d). Of note, eOPN3 is a GPCR and thus does not elicit the instantaneous inhibition seen after photoinhibition using GtACR, thus precluding studies of inhibition in a closed loop paradigm^22^. However, constant inhibition (light) in an open loop configuration decreased food approach (329% latency, see Extended Data Fig.6 e,f) and increased chewing of spaghetti and non-nutritious wooden sticks (wood:642%, spaghetti: 394%, see Fig.6 e-h and supplementary videos 4-6). This suggested that inhibition of VMH^BDNF^ projections to Me5 might directly regulate consummatory actions including movement of the jaw.

**Figure 6.**
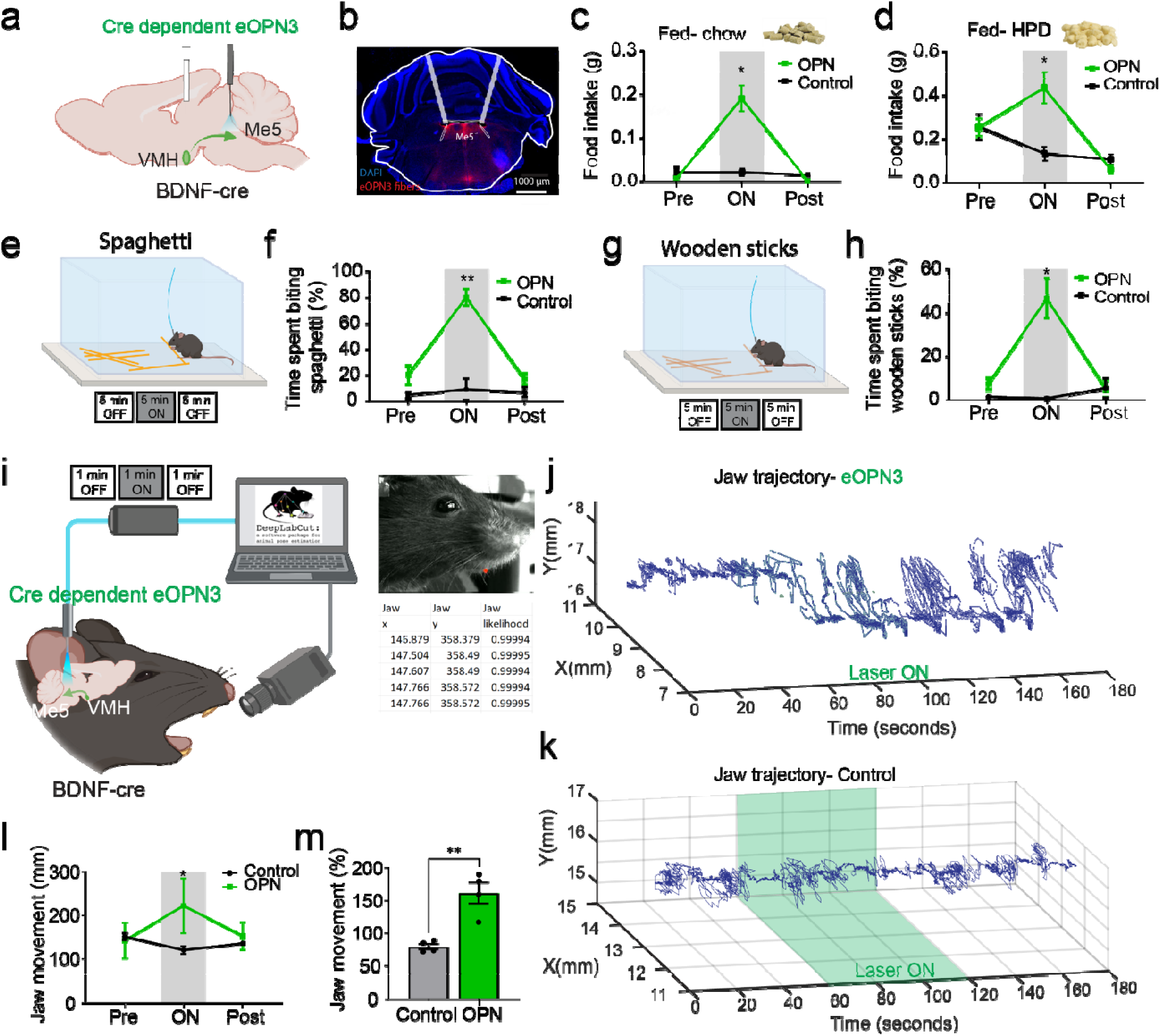
Inhibition of VMH^BDNF^ projections to Me5. **a,** schematic of a sagittal brain with eOPN3 expression in the VMH and fiber placement above Me5. **b**, representative image of eOPN3-Ruby expression and fiber placement above Me5.**c,** food intake of ad lib fed mice tested with acute chow (OPN: 0.19±0.029g, Ctrl: 0.022±0.009g, P=0.0102 Two-way RM ANOVA with Šídák’s multiple comparisons test) and **(d)** ad lib fed tested with HPD (OPN: 0.434±0.072g, Ctrl: 0.132±0.030g, P=0.0301 Two-way RM ANOVA with Šídák’s multiple comparisons test) OPN (n=5) and control mice (n=5). **e,** Schematic of constant optogenetic inhibition during Spaghetti feeding and experimental timeline. **f**, quantification of time spent biting Spaghetti of ad lib fed mice, n=4 (OPN: 80.278±6.215%, Ctrl: 8.739±8.336%, P=0.0019 Two-way RM ANOVA with Šídák’s multiple comparisons test). **g**, Schematic of constant optogenetic inhibition during wood stick biting and experimental timeline. **h,** quantification of time spent biting wood sticks of ad lib fed mice, n=4 (OPN: 46.655±9.098%, Ctrl: 0.450±0.450%, P=0.0432 Two-way RM ANOVA with Šídák’s multiple comparisons test). **i**, schematic of high-speed jaw tracking in head-fixed mice with concurrent optogenetic inhibition and subsequent pose estimation with DeepLabCut. **j,** Representative example of the jaw movements in two dimensions over time of an eOPN3 expressing mouse with the time of laser stimulation indicated in green (60-120s). **k,** Representative example of the jaw movements in two dimensions over time of a control mouse with the time of laser stimulation indicated in green (60-120s). **l**, quantification of absolute jaw movements during laser on and off periods in n=4 control and OPN mice (ON:221.317±63.126mm, Pre: 142.108± 142.108mm, P=0.0047 Two-way RM ANOVA with Šídák’s multiple comparisons test).**m,** comparison of jaw movement between control and OPN mice after normalization to laser off period (OPN: 161.3±16.24% Ctrl: 79.44±4.346%, P=0.0028 Unpaired t-test).

We directly assessed jaw movements in the absence of food or other stimuli in head-fixed mice using a high-speed side camera during optogenetic inhibition of the VMH^BDNF^ projections to Me5. Pose estimation of the jaw position was performed with DeepLabCut as reported in Lauer et al., 2022 (Fig.6i). Photoinhibition of VMH^BDNF^ terminals in Me5 using eOPN3 reproducibly evoked rhythmic jaw movements with a similar time course to that reported in studies of food intake (Figure 6 i-l, supplementary video 7). In contrast, control mice without eOPN3 expression showed infrequent jaw movements (supplementary video 8). Photoinhibition of VMH^BDNF^ terminals in Me5 led to a ∼160% increase of jaw movements during laser inhibition compared to controls (Fig. 6m).

## Discussion

A key function of the brain is to generate adaptive behaviors in response to an array of interoceptive and sensory inputs. A full understanding of how behaviors are controlled in higher organisms will thus require the elucidation of neural circuits linking sensory inputs to motor outputs. In this report we show that BDNF neurons in the VMH convey interoceptive inputs to motor outputs associated with feeding. These neurons receive direct monosynaptic inputs from AgRP and POMC neurons in the Arc, two key interoceptive populations. In vivo imaging studies reveal that the activity of VMH^BDNF^ neurons is inversely correlated with feeding, tuned to energy state and modulated by leptin. Optogenetic activation of VMH^BDNF^ neurons prevents feeding while their inhibition increases food intake. In the absence of food, these actions are directed at inedible objects such as a wood block or sticks suggesting that they drive motor sequences associated with feeding. Inhibition of VMH^BDNF^ projections to Me5, a well-established premotor site elicits rhythmic activation of the jaw muscles that are triggered even in the absence of food. In aggregate, these data show that VMH^BDNF^ neurons directly connect interoceptive inputs from the arcuate nucleus to a premotor site that controls jaw movements and other consummatory actions thus defining a simple subcortical circuit that regulates food consumption (Extended data Fig.6m).

Complex behaviors such as feeding which are context dependent, and thus variable, are considered as mechanistically distinct from a reflex in which a defined stimulus generally results in an invariant response^4^. However, as suggested by Sherrington in 1906^2^, the “transition from reflex action to volitional control is not abrupt and sharp.” Sherrington pointed out that reflexes are also under volitional control noting that the cough reflex (and others) can be “checked, released, or modified in its reaction with such variety and seeming independence of external stimuli by the existence of a spontaneous internal process expressed as will”. This line of reasoning suggests the possibility that even among mammals feeding and other complex behaviors might be controlled in part by simple, reflex like circuits that are in turn modulated by descending inputs from higher centers as was also suggested by Crick and Koch^49^. The possibility that relatively simple, subcortical circuits can regulate feeding is consistent with phylogenetic data from lower vertebrates which can regulate feeding with fidelity despite having only a very small or no cortex^50^. In addition, normal feeding is maintained in decorticate rats and rabbits so long as key diencephalic structures are intact^51,52^. The identification of an Arc→VMH^BDNF^→Me5 circuit reported here is consistent with Sherrington’s hypothesis and by also identifying a premotor node regulating consummatory behaviors potentially provides a framework for establishing a hierarchy among the many previously identified nodes that have been shown to control food intake^53,54^. However, the identification of this circuit does not preclude the possibility that other partially overlapping circuits may also regulate feeding by projecting onto Me5 or premotor neurons in other nuclei. Our finding that VMH^BDNF^ projections to Me5 regulate consummatory behaviors is consistent with lesioning studies in which a knockout of the transcription factor Drg11 led to the ablation of Me5. These mice developed a Me5 lesion and starved to death at weaning due to an inability to consume solid food but survived if provided with liquid nutrient^55^. Our finding of a premotor site regulating consummatory behaviors, thus raises the possibility that other innate behaviors are controlled by premotor sites elsewhere. Tinbergen suggested that key nodes regulating motor patterns are released from inhibition in response to sensory and interoceptive inputs^56^ and our identification of a node regulating food consumption may thus have general implications for how specific behaviors are selected to the exclusion of other competing behaviors ^1^.

These data also show that VMH^BDNF^ neurons are a key functional output of AgRP neurons which play an essential role to drive food intake when an animal’s energy state is low. AgRP levels are suppressed by leptin^57^ and our findings thus suggest that the obesity seen with BDNF, TrkB, leptin, LepR receptor and melanocortin mutations is caused by altered function of the same feeding circuit. While AgRP neuron activation by itself normally drives feeding, concurrent VMH^BDNF^ neuron activation entirely blocks this. AgRP neurons act as interoceptive sensors of changes in energy state and drive all aspects of hunger^58,59^ including both appetitive behaviors such as food seeking and consummatory behaviors. However, we find that AgRP inputs to VMH^BDNF^ neurons increase only the consummatory phase of feeding but not food approach, suggesting that different projection sites of these neurons regulate the motivational component. This is consistent with a hierarchical architecture for innate behaviors as proposed by Craig^60^, Lorenz^3^ and especially Tinbergen^1^ in which appetitive and consummatory behaviors are controlled by different sites and that the initiation of the consummatory phase only begins when the appetitive phase is completed and food is in proximity. Moreover, AgRP neuron projections to the VMH do not drive the consummatory motor sequences by themselves in the absence of food, suggesting that additional signals are necessary to ‘release’ the motor programs for mastication. As suggested by Tinbergen, such inhibition could be sensory information derived from potential food sources signaling that food is in proximity. The possibility that AgRP inputs to VMH^BDNF^ neurons are not sufficient to activate consumption is analogous to Tinbergen’s experiment in which honey bees only land on colorful artificial flowers when they are paired with appropriate odours^56^.

The data also show that VMH^BDNF^ neurons are a critical node downstream of leptin signaling as shown by our findings that leptin increases the gain of these neurons using fiber photometry and DtA ablation of VMH^BDNF^ neurons significantly decreases leptin’s effect on food intake. Retrograde viral tracing from VMH^BDNF^ neurons identified inputs from AgRP and POMC in the arcuate nucleus, the site of a circumventricular organ, that convey the leptin signal to VMH^BDNF^ neurons which themselves do not express leptin receptors. Modulation of VMH^BDNF^ neural activity entirely blocks the effect of POMC or AgRP neural activation on feeding confirming that they are a key functional target of these neurons. This finding is in line with prior studies reporting the presence of POMC and AgRP fibers and terminals in the VMH^34,61^. Consistent with this, VMH^BDNF^ neurons co-express MC4R and NPY5R suggesting that they integrate both melanocortin and NPY feeding signals^34^ and are a key integrator. Previous studies^62^ also suggested the possibility that VMH^BDNF^ were downstream of MC4R signaling. For example, Xu et al reported that BDNF injections can ameliorate the hyperphagic phenotype of A^Y^ mice and that BDNF expression in the VMH is regulated by melanocortin signaling. We found that the activity of VMH^BDNF^ neurons can be modulated by leptin and by additional factors because ablating VMH^BDNF^ neurons increases weight in ob/ob animals lacking leptin. The data presented here are thus consistent with the possibility that both leptin dependent and leptin independent signals regulate the activity of VMH^BDNF^ neurons. A leptin independent effect is also consistent with a prior report showing that ob/ob mice receiving a six month infusion of leptin while on a high fat diet end up at a similar weight to wild type animals fed the same diet^63^. The fact that leptin levels were fixed in these ob/ob mice suggests that another signal(s) besides leptin was limiting the weight of this group. Another study also suggested the existence of additional signals besides leptin^64^ and it will now be important to identify the putative leptin independent signals which might or might not be mediated by melanocortins.

Our findings also suggest that ablation of VMH^BDNF^ neurons accounts for the massive obesity associated with lesions of the VMH in rodents^65,66^ and humans^67,68^. This effect is not recapitulated by a knockout of Sf1, a canonical marker expressed in most VMH neurons, or a knockout of BDNF in Sf1 neurons^17^ suggesting that the VMH^BDNF^ neurons are distinct from Sf1 expressing neurons there. A postnatal knockout of Sf1 led only to mild obesity in mice fed a high fat diet as a result of impaired thermogenesis with little or no effect on food intake^69^ and there was only a minimal effect on food intake or weight of chow fed mice. We find only limited overlap between BDNF and Sf1 neurons which mark nearly all of the other neurons in this nucleus (∼6% of SF1 neurons also express BDNF according to single cell RNA sequencing)^15^. These data thus suggest that it is the subpopulation of BDNF neurons in VMH that accounts for the hyperphagia and massive obesity that follows a VMH lesion. These data also suggest that VMH^BDNF^ neurons contribute significantly to the obesity associated with BDNF and TrkB mutations. Further support for this conclusion was provided in a recent study that found an interaction of BDNF and astrocytes in the VMH that is essential for the suppression of weight ^70^. However, this does not exclude the possibility that other sites might also contribute to the obesity associated with a BDNF or TrkB mutation. For example, while they do not regulate feeding, BDNF expressing neurons in the paraventricular nucleus have been shown regulate energy expenditure with effects on thermogenesis and adipose tissue innervation^71,72^.

The data further suggest that VMH^BDNF^ neurons play a role in defining a set point for body weight once obesity develops. Large scale longitudinal studies have shown that body weight is remarkably stable over the course of a year (or longer) not just among lean but also among obese individuals. For example, body weight typically returns to one’s baseline even after periods associated with increased intake of calorie dense palatable meals such as Christmas and Golden Week^73^. This has suggested that weight and adipose tissue mass are under homeostatic control even among obese individuals or when highly palatable food is available. Similarly, when provided with highly palatable food, C57/Bl6 mice initially binge eat but as they approach a new, higher stable weight their food intake decreases. Our observation that VMH^BDNF^ neurons show c-fos activation and increased GCaMP transients in DIO animals (Fig.1a and 3d-j suggests that these neurons are tuned to an animal’s energy state and that these neurons act to prevent further weight gain and the development of an even more severe obese phenotype in mice fed HPD. Consistent with this ablating VMH^BDNF^ neurons increases body weight in both DIO and ob/ob mice. These data combined with the results of optogenetic manipulations suggest that these neurons act to maintain the set point even after obesity develops.

In summary, these data show that VMH^BDNF^ neurons are a key element of a simple circuit that regulates food intake and body weight by directly connecting interoceptive inputs to motor outputs controlling consummatory behaviors and jaw movements. These results now provide a framework for studying the neural mechanisms by which higher centers may modulate feeding via actions on the Arc→VMH^BDNF^→Me5 or possibly other minimal circuits. These findings thus have important implications for the regulation of feeding and potentially other complex motivational behaviors.

## Supporting information

video1_opto_wood_block

video2_photometry_pellet

video3_photometry_rejection

video4_spaghetti_laserOFF_4x

video5_spaghetti_laserON_4x

video6_woodensticks_laserON_4x

video7_OPN_jaw_5x

video8_OPN_control_jaw_5x

## Methods

### Mice

All animal care and experimentation were ethically performed according to procedures approved by the Institutional Animal Care and Use Committee at Rockefeller University. Male mice were single housed in a 12-h light/12-h dark cycle with ad libitum access to regular chow and water except in fasting and DIO studies where either a HFD with 45 kcal%fat (4.7kcal/g) or a HPD with 42 kcal%fat and high sucrose content (4.5kcal/g) (TD.88137 Envigo) were provided. We used male ob/ob (B6.Cg-Lep^ob^/J; 000632, Jackson Laboratory; or bred in-house), Rosa26^fsTRAP^ (B6.129S4-Gt(ROSA)26Sor^tm1(CAG-EGFP/Rpl10a,-birA)Wtp^/J; 022367, Jackson Laboratory), AgRP-cre (AgRP^tm1(cre)Lowl^/J; 012899, Jackson Laboratory), Flp reporter mice (RCF-tdTomato, B6.Cg-Gt(ROSA)26Sortm65.2(CAG-tdTomato)Hze/J; 032864, Jackson Laboratory) crossed to cre reporter mice (Rosa26^fsTRAP^). BDNF-IRES-Cre mice were provided by W. Shen (Shanghai Institute of Technology). POMC-dre were provided by J. Bruning (Max Planck Institute for Metabolism Research). For retrograde tracing from motor neurons in Mo5 Chat cre mice (B6.129S-Chattm1(cre)Lowl/MwarJ; 031661, Jackson Laboratory) were crossed to Helper RabV mice (B6;129P2-Gt(ROSA)26Sortm1(CAG-RABVgp4,-TVA)Arenk/J; 024708, Jackson Laboratory). All mouse lines are in a WT (C57BL/6J) background. For brain surgeries, male mice of at least 8 weeks of age were anesthetized with isoflurane and placed into a stereotaxic frame (David Kopf Instruments), a craniotomy was performed, and a borosilicate glass pipette was used to inject viral vectors. For VMH injections: three injections (each 50LJnL) were made into each hemisphere (bregma: −1.36LJmm, midline: ±0.35 mm, from brain surface: 5.70 mm, 5.60 and 5.50 mm). For injections into the Arc 50nl were injected into: bregma: −1.45LJmm, midline: ±0.45 mm, from brain surface: 5.70 mm, 5.60 and 5.50 mm, For injections into Mo5 75nl were injected into: bregma: −5.20mm, midline:±1.5mm, from brain surface: 4.60 and 4.50mm. For Me5 injections from bregma: −5.4LJmm, midline: ±0.9 mm, from brain surface: 4.5, 4.0, 3.5mm.

### Reagents

leptin was diluted in sterile saline (3mg/kg) and injected intraperitoneally. For photometry recording mice were overnight fasted and injected with either saline or leptin 2h before recordings. All mice received saline and leptin injections in alternating order. For acute food intake experiments, leptin or saline was injected 2h before onset of the dark period and all mice received saline and leptin injections in a cross-over design.

### Viruses

Cre dependent neuronal ablation was performed by injection of AAV1-mCherry-flex-dtA (UNC Vector Core)^20^. To target expression of the calcium activity indicator GCaMP6s to VMH^BDNF^ neurons, we used an AAV vector carrying a double floxed GCaMP6s construct (AAV5-Syn-Flex-GCaMP6s-WPRE-SV40, Addgene) ^74^. For optogenetic manipulations, a somatic targeting GtACR (AAV5-hSyn1-SIO-stGtACR1-FusionRed) ^22^ or ChR (AAV5-EF1a-double floxed-hChR2(H134R)-EYFP-WPRE-HGHpA) were used (both Addgene). For long-term silencing a cre dependent TelC AAV (AAV5-hSyn-FLEX-TeLC-P2A-dTomato, Addgene) was used and for long-term activation a cre dependent NaChBac (AAV-Syn-DIO-NaChBac-dTomato) and a Dre dependent NaChBac (AAV5-hSyn-roxSTOProx-NaChBac-dTomato, HHMI-Janelia Research Campus) were used. For retrograde tracing a combination of two helper AAVs (AAV1-TREtight-mTagBFP2-B19G and AAV1-syn-FLEX-splitTVA-EGFP-tTA, both Addgene) and pseudotyped rabies (EnvA G-Deleted Rabies-mCherry, Salk Institute) were injected, for retrograde labelling from Mo5 a G-Deleted Rabies-H2B-mCherry (Salk viral core) was used and for anatomical tracing from Me5 a retrograde mCherry construct (pAAV-hSyn-DIO-hM4D(Gi)-mCherry, Addgene) was used. For projection activation we used an AAV encoding eOPN3 (AAV-hSyn1-SIO-eOPN3-mScarlet-WPRE, Addgene)^22^ and for labelling projections with ChR we used a retro-AAV (AAV-EF1a-double floxed-hChR2(H134R)-mCherry-WPRE-HGHpA). Anterograde labelling was done with AAV1 encoding cre (AAV-hSyn-Cre-WPRE-hGH, Addgene) and flp (AAV-EF1a-Flpo, Addgene). For ‘cre-out’ experiments a (AAV-Ef1a-DO-ChETA-EYFP-WPRE-pA, Addgene) was used.

### Immunofluorescence

C-Fos staining after DIO: BDNF-cre mice were crossed to Rosa26fsTRAP to express eGFP in a cre dependent manner in BDNF neurons. Mice were fed a HFD whilst littermate control mice were fed chow. After 16 weeks mice were transcardially perfused with 4% PFA and the brains were post-fixed for 1 day in 4% PFA. Brains were then placed in 30% Sucrose in PBS until precipitation and frozen and coated in O.C.T. for cryosectioning. Cryosections (50μm) were cut using a Leica cryostat (CM1950). Brain sections were washed in PBS with 0.1% Triton X-100 (PBST, pH 7.4) and blocked in 3% normal goat/donkey serum (Jackson ImmunoResearch Laboratories) and 2% BSA (Sigma) in PBST for 2 h. Slides were then incubated overnight at room temperature in primary antibody. After washing in PBST, sections were incubated in fluorescein-conjugated goat IgG. Primary antibodies used and their dilutions: rabbit anti-FOS (1:1,000; mAb 2250S, Cell Signaling), chicken anti-GFP (1:1,000, ab 13970, Abcam). Secondary antibodies conjugated with Alexa-594, and Alexa-488 were purchased from Invitrogen. Brain sections were mounted onto SuperFrost (Fisher Scientific 22-034-980) slides and then visualized with an inverted Zeiss LSM 780 laser scanning confocal microscope with a ×10 or ×20 lens. Images were imported to Fiji for further analysis and to count cells. To quantify the number of stained cells, brain slides were imaged under a ×20 objective. For advillin staining: same as above but with rabbit-anti advillin (1:500, NBP2-92263, Novus Biologicals) for primary and Alexa 647 donkey anti-rabbit (1:500, ab150075, Abcam) for secondary antibody. For anterograde tracing: Brains were processed as described above and tdtomato/Ruby and GFP were amplified with rabbit anti-RFP (1:1000, 600-401-379, Rockland) and chicken anti-GFP (1:1,000, ab 13970, Abcam) as primary antibodies and secondary antibodies conjugated with Alexa-594, and Alexa-488 were purchased from Invitrogen.

### In situ hybridization

Mice were briefly transcardially perfused with RNase-free PBS to remove blood. Brains were then quickly collected, frozen in OCT, and stored at −80LJ°C until sectioning by cryostat (15μm sections) and attached on Superfrost Plus Adhesion Slides (Thermo Fisher). RNAscope® Fluorescent Multiplex assay (Advanced Cell Diagnostics Bio) was then performed using the RNAscope system. Probes for the following mRNAs were used (all from ACDBio): mm-BDNF (cat no. 424821) and eGFP (cat no. 400281), VGlut2 (cat no. 319171), RabV (cat no. 456781), AgRP (cat no. 400711), POMC (cat no. 314081), MC4R (cat no. 319181-C2) and NPY5R (cat no. 589811), LepR (cat no. 402731). In brief, RNAscope was used as per the manufacturer’s protocol. Briefly, a hydrophobic barrier was created using Immedge Hydrophobic Barrier Pen (Vector Laboratories). Pre-treatment was done by serial submersion of the slides in 1× PBS, 50% EtOH, 70% EtOH and twice 100% EtOH for 2 min each, at room temperature. Probe hybridization was achieved by incubation of 35 μl mRNA target probes for 2 h at 40LJ°C using a HyBez oven. The signal was amplified by subsequent incubation of Amp-1, Amp-2, Amp-3 and Amp-4 one drop each for 30, 15, 30 and 15 min, respectively, at 40LJ°C using a HyBez oven. Each incubation step was followed by two 2-min washes using RNAscope washing buffer. Nucleic acids were stained using DAPI Fluoromount-G (SouthernBiotech) mounting medium before coverslipping. Slides were visualized with an inverted Zeiss LSM 780 laser scanning confocal microscope using a ×20 or ×40 lens. Images were imported to Fiji for further analysis.

### Long term body weight and food intake measures

Single housed mice were measured weekly to assess body weight and food intake. Whole-body composition was measured using Nuclear Magnetic Resonance Relaxometry (EchoMRI) at the end of the 16 weeks period.

### Optogenetics

After injection of AAVs encoding either ChR or GtACR, we bilaterally implanted 200μm fiber optic cannulas (Thorlabs) in BDNF-cre mice and control mice (cre negative littermates). For VMH targeting, implants were angled at 15° and placed at bregma: −1.36LJmm, midline: ±1.85LJmm, from brain surface: 5.25 mm, for brainstem targeting implants were angled at 15° and placed at bregma: −5.4LJmm, midline: ±1.75LJmm, from brain surface: 2.6 mm for Me5, −6.3LJmm, midline: ±2.15LJmm, from brain surface: 5.5 mm for LPGi, −5.7LJmm, midline: ±2.6LJmm, from brain surface: 4.7 mm for PCRT and subsequently fixed by dental cement (C&B metabond). After a minimum of 3 weeks expression time, mice were handled and habituated to tethering with optical fibers. A constant 473 nm laser (OEM Lasers/OptoEngine) was used for optogenetic inhibition with GtACR and pulsed at 2Hz (5ms) for optoactivation with ChR. For inhibition with OPN, a 532nm laser at 10Hz was used. Lasers were connected to bifurcated optical fibers (Thorlabs) with an output of ∼ 2-5mW at the implant., for AgRP projection stimulation the laser power was reduced to 1-2mW. Food intake studies were done in home cage like arenas during the light phase without bedding unless otherwise stated.

<u>For acute food intake experiments</u> with optogenetic activation or inhibition, mice were habituated to the arena for 10min without food present. Then consumption of a single food pellet was measured every 30min for 5 repetitions with only the second repetition being paired with optogenetic activation or inhibition. For <u>open loop and closed loop feeding experiments</u>, a single chow pellet was fixed to the middle of a home cage style arena with fun-tak (Loctite). Food intake was assessed every 5min in 3 repetitions with only the second repetition being paired with optogenetic inhibition. Inhibition was either 5min constant laser (open loop) or triggered (closed loop) by real time video tracking (Noldus, Ethovision) whenever the head of the mouse was within a 3 cm radius of the pellet. For modification with bedding present, the same open loop setup was used but with corn cob bedding covering the floor. For wood block trials, the chow pellet was replaced by a wood block that was fixed with fun-tak. Time spent biting the wood block was manually assessed and quantified by scoring of video recordings.

<u>Liquid diet experiments</u> were done with 20µl Ensure Vanilla pipetted onto the bare floor of a cage in 3 repetitions without light activation, followed by 3 repetitions with light activation and 3 repetitions without light activation. For quantification purposes, experiments were video recorded and latencies from Ensure delivery to full consumption were scored and averaged over the 3 repetitions.

<u>Operant conditioning</u> was performed in a home cage style arena with two capacitive touch plates mounted on opposing sides. Both touch plates were connected to an Arduino to register numbers of touches and one randomly assigned side would trigger cessation of constant 2Hz laser (for ChR) for 3 seconds or activation of a constant laser for 3 sec (for GtACR). Trials lasted for 1h.

<u>Conditioned Flavor preference assays</u> were performed as previously described ^59^. Briefly, mice were habituated overnight to orange and strawberry flavored sugar-free Juicy Gels (Hunt’s). Initial preference was assessed in a 30 min session without any light application. The preferred flavor was then paired with light exposure for ChR mice or the less preferred flavor paired with light exposure for GtACR mice. Conditioning was repeated daily for 3 days and consisted of one light exposure session where after 5 min light exposure started and lasted for 25min whilst the paired gel was presented and a 30min session with the non-paired gel without any light exposure. A 15 min test session where both gels were available was performed on the day after conditioning ended.

<u>Spaghetti and wooden stick experiments:</u> 5 spaghetti sticks or wooden sticks of similar length were distributed equally in an empty home cage. Control and eOPN3 mice were given 5 min baseline exploration, 5 min with laser inhibition and 5 min without laser with the spaghetti or sticks present. A side and overhead camera were used to quantify the time spent chewing.

### Head-fixed Jaw tracking

Mice for head-fixed experiments had during implant surgery also a small metal bar fixed to their scull with dental cements. After a minimum of 3 weeks recovery, mice were habituated to being head fixed in a custom head-fixation set up. This set up consists of a side camera (Basler a2A1920-160umPRO-ace 2) and laser source controlled and synchronized by Bonsai^75^. Frames were acquired at 100Hz at 722 × 878 pixel size. Optogenetic inhibition trials consisted of 1min with laser, 1min on and 1 min off. Jaw pose was subsequently estimated with DeepLabCut^48^.

### Fiber photometry

After an injection of AAV encoding cre dependent GCaMP6s into the VMH of male BDNF-cre mice, a unilateral 400μm fiber optic cannula was implanted as described for optogenetics. After a minimum of 4 weeks expression time, mice were habituated to tethering and a home cage style arena.

Data was collected with a Fiber Photometry system by Tucker-Davis Technologies (RZ5P, Synapse) and Doric components and recordings were synched to video recordings in Ethovision via TTL triggering. A 465nm and isosbestic 405nm LED (Doric) were reflected into a dual fluorescence Mini Cube (Doric) before entering the recording fiber that connects to the implant. Recording fibers were photobleached overnight before recordings to minimize autofluorescence. GCaMP6s fluorescence was collected as calcium dependent signal (525nm) and isosbestic control (430nm) with a femtowatt photoreceivers (Newport, 2151) and a lock-in amplifier via the RZ5P at a 1kHz sampling rate.

Mice were allowed to habituate for 30min at the start of each recording session before any items were introduced into the arena. Feeding bouts were manually assessed and scored from video recordings when mice were given single pellets of chow or 20mg sucrose treat pellets (Bio-Serv). Food interaction without consumption instances were defined as approach within a 2cm radius around a chow pellet without subsequent consumption. To measure the effect of different energy states, the same mice underwent the same standardized recording procedure at lean ad lib chow fed, overnight fasted injected with saline, overnight fasted injected with leptin and 4 weeks DIO. The order between lean, fasted saline, and fasted leptin were randomized to avoid any order effects.

Analysis was performed with a script written in Matlab based on a previously published method and code^76^. Bleaching and movement artefacts were removed by applying a polynomial least-square fit to the 405nm signal adjusting it to the 465nm trace (405_fitted_) to then calculate the GCaMP signal as % ΔF/F = (465_signal_-405_fitted_)/405_fitted_. Traces were filtered with a moving average filter and down sampled by a factor 20. Three trials per mouse were averaged to derive data for peri-event plots and analysis of maximum and minimum signals.

## Quantification and Statistic

Microscope images were analyzed and quantified in ImageJ/Fiji. Photometry recordings were processed and analyzed with MatLab (Mathworks). Statistical analysis was performed in GraphPad Prism. All tests are two-sided and results are displayed as mean±sem with statistical details in the figure legends including definition of n and significance. Significance was defined as p<0.05. Significance annotations are: ∗p<0.05, ∗∗p<0.01, ∗∗∗p<0.001, ∗∗∗∗p<0.0001. Mice were randomized into control or treatment groups. Control mice were age-matched littermate controls where possible. Graphs were produced using GraphPad Prism and Adobe illustrator with schematic illustrations prepared in BioRender.

## Data availability

The data that support the findings of this study are available from the corresponding authors upon reasonable request. Requests for reagents should be directed to J.M.F.

## Code availability

Analysis code for fiber photometry, Arduino code for operant conditioning, DeepLabCut code for jaw pose and Bonsai code for head-fixation behavior are available upon request.

## Acknowledgements

We thank all members of the Friedman lab for discussions and comments on the project, especially Anoj Ilanges. K. Hedbacker for breeding ob/ob mice, Rockefeller Bio-Imaging Resource Center (BIRC) for microscopy and Rockefeller Precision Instrumentation Technologies (PIT) for fabrication of head bars and help with the head-fixed setup. We thank Dr Bruning (Max Planck Institute) for the provision of POMC Dre mice. We thank Robert Smith for assistance with Arduino and DeepLabCut implementation. This work was supported by the JPB Foundation. C. K. was supported by a fellowship from the Leon-Levy Foundation.

## Author contributions

C.K designed and performed all experiments. J.F. and C.K wrote the manuscript. Z.K. designed and performed TrkB mutant experiments. J.I and K.P. assisted C.K with experiments.

## Supplementary information

Supplementary Video 1: Optogenetic inhibiting of VMH^BDNF^ neurons when a mouse is near a wood block.

Supplementary Video 2: Photometry recording of a mouse approaching and consuming a treat pellet.

Supplementary Video 3: Photometry recording of a mouse approaching and not consuming a chow pellet.

Supplementary Video 4: Representative recording of OPN mouse interacting with Spaghetti without laser (4x speed).

Supplementary Video 5: Representative recording of OPN mouse interacting with Spaghetti during VMH^BDNF^ to Me5 inhibition (4x speed).

Supplementary Video 6: Representative recording of OPN mice interacting with wooden sticks during VMH^BDNF^ to Me5 inhibition (4x speed).

Supplementary Video 7: Jaw movements and tracking during VMH^BDNF^ neuron to Me5 inhibition (speed 5x).

Supplementary Video 8: Jaw movements and tracking in a control mouse (speed 5x).

## Data Availability

The datasets generated and analyzed during the current study are available from the corresponding author on reasonable request.

## Code Availability

The code generated during the current study are available from the corresponding author on reasonable request.

## Extended data figure legends

**Extended data figure 1: Overlap of VMH^BDNF^ neurons with Vglut2 and Sf1 and effect of TrkB inhibition**

**a,** representative example of coronal sections (n=3 mice) through the VMH with in situ hybridization for BDNF (magenta) and Sf1 (white) and DAPI (blue). **b,** quantification of overlap with SF1 and BDNF in dorsomedial and central VMH. **c**, high magnification examples of overlap (indicated by white arrows) of boxed areas from a. **d,** representative example of coronal sections (n=3 mice) through the VMH with in situ hybridization for BDNF (magenta) and Vglut2 (green) and DAPI (blue). **e,** high magnification examples of overlap (indicated by white arrows) of boxed areas from d. Right, quantification of Glut2 expression in VMH^BDNF^ neurons of 3 mice.

**Extended data figure 2: Inhibition of VMH^BDNF^ neurons in the DIO state and VMH^BDNF^ neuron effects on valence and social interaction**

**a,** schematic of the HPD induced obese state. **b,** changes in body after DIO. **c,** quantification of food intake before, during after optoinhibiton in DIO mice expressing GTaCR (n=6) and control mice (n=6) given chow: for GtACR 0.155±0.023g vs 0.013±0.003 for controls, P= <0.0001, Two-way RM ANOVA with Šídák’s multiple comparisons test, and **d,** HPD: for GtACR 0.358±0.006g vs 0.015±0.006g for controls, P= 0.0123, Two-way RM ANOVA with Šídák’s multiple comparisons test. **e,** quantification of chow intake (0.198±0.037g in lean vs 0.066±0.016g in DIO, P=0.0019, Paired t-test), and in **f,** HPD intake (0.541±0.050g in lean vs 0.1317±0.048g in DIO, P<0.0001, Paired t-test), without optoinhibition in all (n=12) mice when feed chow chronically and after DIO. **g**, quantification of the optoinhibition evoked feeding (difference between food intake with and without laser), 0.168±0.067g in lean vs 0.313±0.072g in DIO, P=0.0146, Two-way RM ANOVA with Šídák’s multiple comparisons test. **h,** Time course of body weight before and during 1-NM PP1 treatments in control (n=4 each) and TrkB mutant mice (body weight change: 16.9±1.17% mutant vs 0.5±2.18% in controls, P=0.0258, Two-way RM ANOVA with Šídák’s multiple comparisons test). **i,** time course of food intake and **j,** quantification of average food intake before and during 1-NM PP1 treatment (21.17±1.18kcal mutant vs in control 16.67±2.24kcal, P <0.0001, Two-way RM ANOVA with Šídák’s multiple comparisons test). **k,** schematic of set up for touch activated self-inhibition with an active site that controls the activity of a laser. **l,** Quantification of the number of touches at the active and inactive site for ChR (n=6) and control mice (n=6) during self-inhibition (Two-way RM ANOVA with Šídák’s multiple comparisons test, ChR: P=0.9476, Control: P=0.9973). **m,** quantification of the number of touches at the active and inactive site for GtACR (n=6) and control mice (n=6) during self-inhibition (Two-way RM ANOVA with Šídák’s multiple comparisons test, GtACR: P=0.6916, Control: P=0.9883). **n,** schematic of conditioned flavour preference assay. **o,** Quantification of flavour preference at the initial preference test (Pre) and conditioned preference test (post) of conditioned flavour avoidance in ChR (n=6) and control mice (n=6) (Two-way RM ANOVA with Šídák’s multiple comparisons test, ChR: P=0.1296, Control: P=0.9067) and quantification of the change in preference (Welch’s t test, P= 0.2768). **p,** left, Quantification of flavour preference at the initial preference test (Pre) and conditioned preference test (post) of conditioned flavour preference in GtACR (n=6) and control mice (n=6) (Two-way RM ANOVA with Šídák’s multiple comparisons test, GtACR: P=0.1481, Control: P=0.1957) and right, quantification of the change in preference (Welch’s t test, P= 0.6814).**q,** schematic of social interaction assay with optoinibtion. **r**, quantification of timer spent in social interactions with and with laser for mice (n=6) expressing GtACR and control mice (n=6). GtACR: 4.414±0.833% and Ctrl: 18.271±1.749%, P=0.0005, Two-way RM ANOVA with Šídák’s multiple comparisons test.

**Extended data figure 3: Detector unit counts of photometry mice in fed, fasted and DIO state**

**a,** 200s long example traces in detector units (not normalized) during empty cage exploration and **b,** comparison of the median thereof. Fasted: 773.6±153.0, Fed 792.1±136.2, HPD 850.8±121.6, P= 0.4687, Repeated measures one-way ANOVA.

**Extended data figure 4 Leptin engages VMH^BDNF^ neurons to suppress feeding**

**a,** schematic of experimental design. **b,** average photometry trace (±sem) of mice injected with saline (purple) or leptin (green) (n=6 mice) aligned to a bout of treat pellet consumption. **c,** comparison of the average photometry signal during food consumption between 6 mice treated with leptin and saline (−4.792± 0.5139% saline injected vs in −3.550± 0.6081% leptin injected, P= 0.0392, Paired t-test). **d,** schematic of experimental design. **e,** food intake in 24h after a leptin or saline ip injection in Dta and control mice (n=8 each group) (DtA: p=0.0031, Ctrl: p<0.0001, 2-way ANOVA with Holm-Šídák’s multiple comparisons test). **f,** quantification of the feeding suppressive effect of leptin by normalizing to the saline injected food intake (DtA: 11.48%± 2.895, Ctrl: 23.64%±3.513, p= 0.0187, Welch’s t-test). **g**, weekly time course of body weight and **h,** calorie intake of chow fed ob/ob mice crossed to BDNF cre (n=7) and ob/ob control mice (n=7) injected with DtA (BW at week 12: DtA: 124.3%± 2.265, ctrl:112.8%± 3.682, p= 0.0248, Welch’s t-test). **i,** quantification of food intake before and after DtA ablation (obob DtA:16.88±1.362kcal at baseline vs 18.24±1.019 after ablation, P=0.036, Two-way RM ANOVA with Šídák’s multiple comparisons test. **j,** schematic of G-deleted pseudotyped rabies and helper AAV expression. **k,** representative example of a coronal Arc section with in situ hybridization for POMC (white), Rabies (red) and LepRb (green), right, high magnification examples of overlap. **l,** quantification of average co-expression of Rabies labeled POMC neurons with LepRb**. m,** representative example of a coronal Arc section with in situ hybridization for AgRP (white), Rabies (red) and LepRb (green), right, high magnification examples of overlap. **n**, quantification of average co-expression of Rabies labeled AgRP neurons with LepRb.

**Extended data figure 5: VMH^BDNF^ neurons are downstream of melanocortin signaling**

**a,** representative example of a coronal VMH section with in situ hybridization for BDNF (white), MC4R (red) and NPY5R (green), middle, high magnification examples of overlap. **b,** quantification of average co-expression of VMH^BDNF^ neurons with MC4R and NPY5R. **c,** schematic of retrograde ChR expression in Arc and optic fiber placement in VMH and **d**, schematic of open loop optostimulation for food approach assessment. **e,** quantification of latency to reach a food pellet of AgRP cre mice injected with retro ChR into the VMH (n=4) and control mice (n=4). **f**, schematic of G-deleted pseudotyped rabies and helper AAV injection. **g,** representative example of a coronal Arc section with in situ hybridization for MC4R (white) and Rabies (red), right, high magnification examples of overlap and quantification.

**Extended data figure 6: VMH^BDNF^ neuron projections to brainstem drive solid food intake a**, schematic of open loop optogenetic activation with ChR and experimental timeline. **b,** latency to approach the chow pellet in ChR and Ctrl mice. ChR: 15.86±3.06s, Control: 14.43±2.59s, P=0.9799, Two-way RM ANOVA with Šídák’s multiple comparisons test **c**, schematic of open loop optogentic activation and experimental timeline for licking measurements. **d,** quantification of time spent licking of ChR (n=7) and control mice (n=7) given 50µl liquid diet in an open loop inhibition setup. ChR: 43.71±10.84s, Control: 37.67±13.93s, P=0.9822, Two-way RM ANOVA with Šídák’s multiple comparisons test. **e**, schematic of open loop optostimulation for food approach assessment. **f,** quantification of latency to reach a food pellet of BDNF cre mice injected with eOPN3 into VMH and fibers implanted above Me5 (n=4) and control mice (n=4). **g,** schematic of a ‘cre out’ Cheta YFP virus injection into VMH of BDNF cre mice, labelling non-BDNF neurons. **h,** representative image of a coronal section of the injection site. **i,** representative image of projections in Barrington nucleus and surrounding areas and high magnification of area in dotted square. Me5 was marked by autofluorescence of neurons in this nucleus. **j**, schematic of virus injection into the CeA and VMH of Cre and Flp reporter expressing mice. **k**, representative example of n=3 mice of brainstem premotor areas (mesencephalic) and **l,** high magnification of AAV1 labelled neurons. **m**, schematic of a proposed circuit for energy state driven food consumption behavior.

