## extended data figures for "A Subcortical Feeding Circuit Linking Interoception to Jaw movement"

**Extended data figure 1**
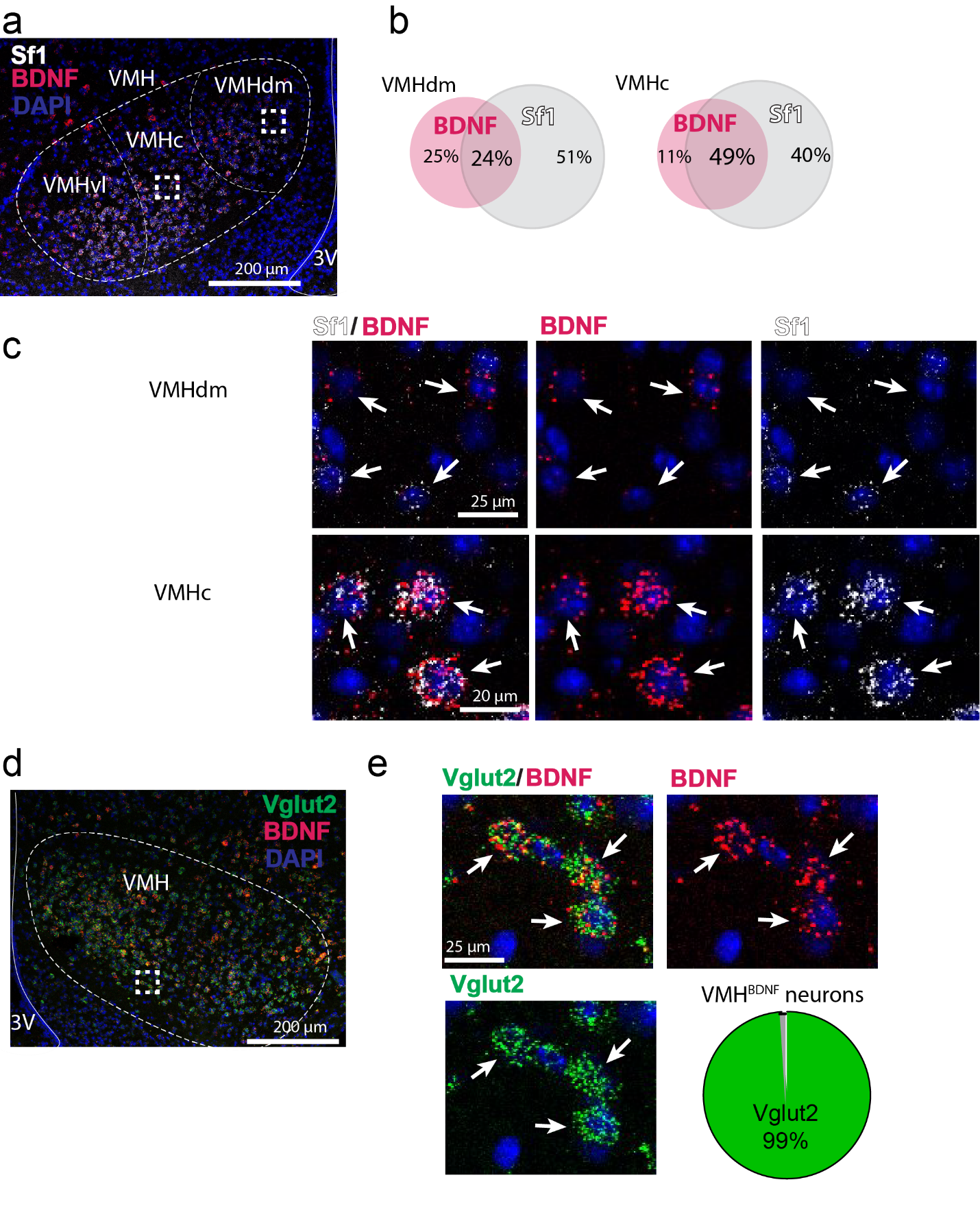

**Extended data figure 2**
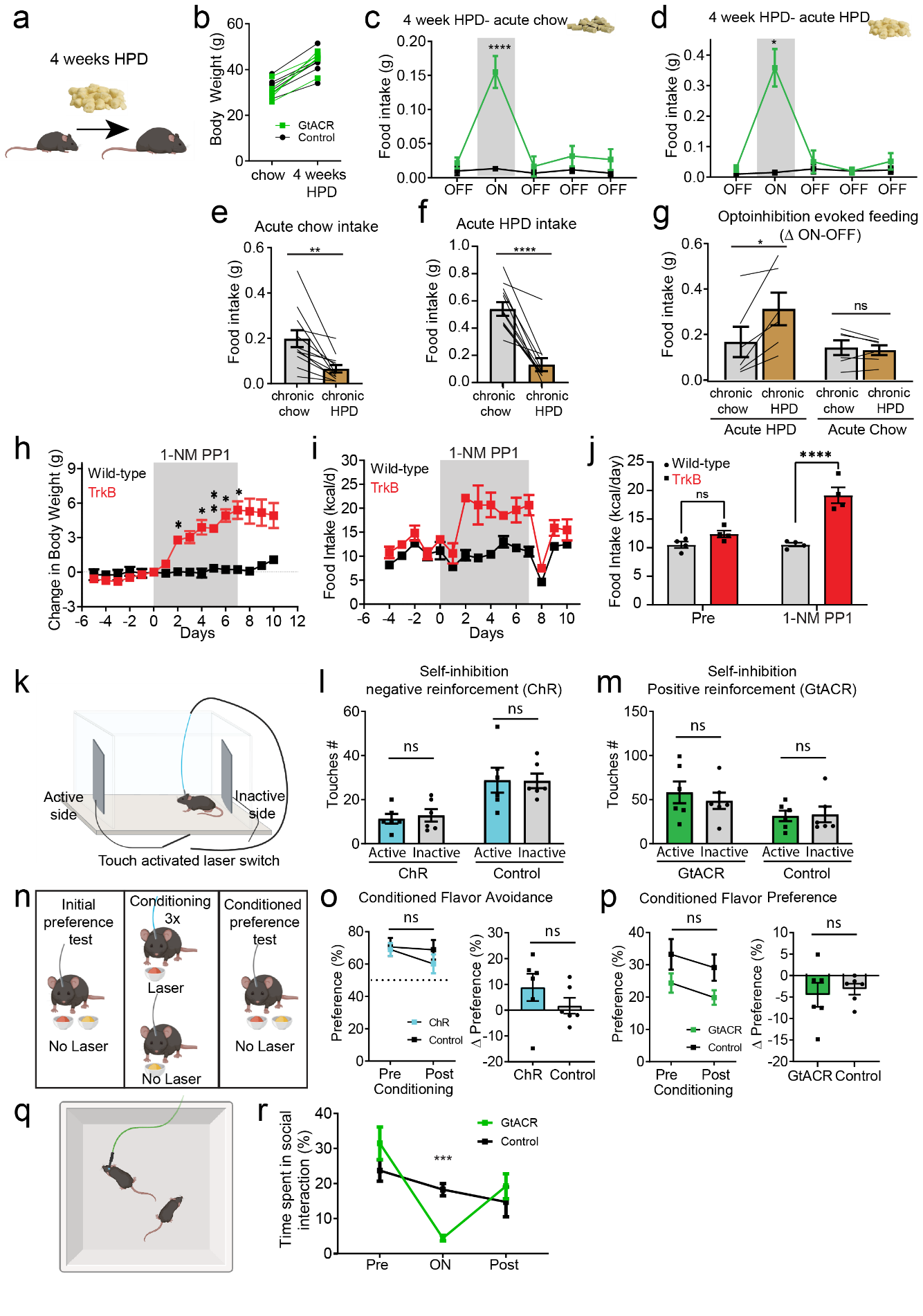

**Extended data figure 3**

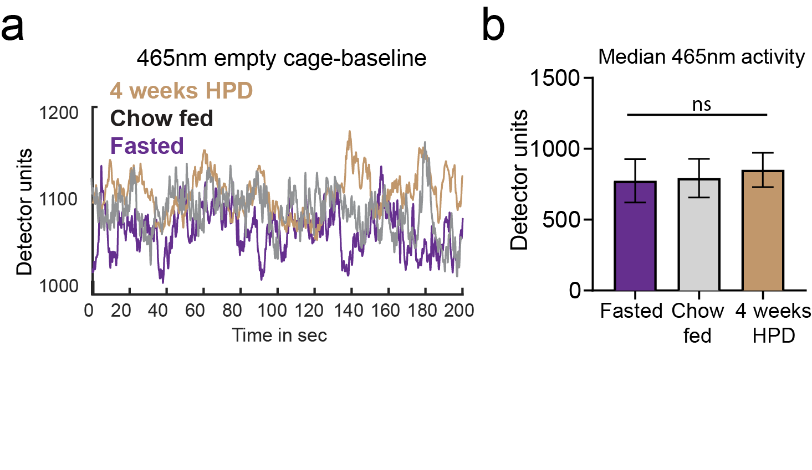

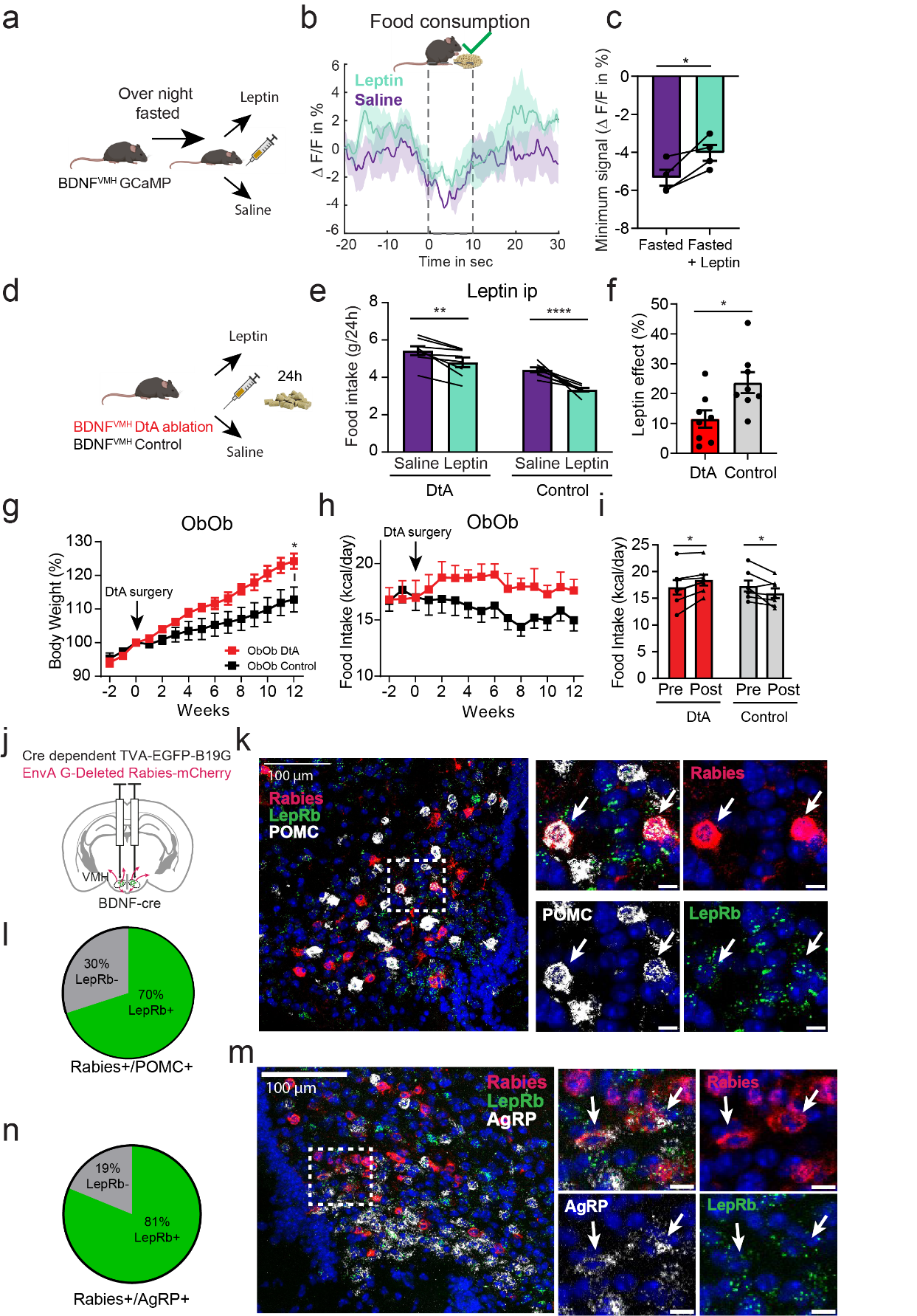
**Extended data figure 4**

**Extended data figure 5**

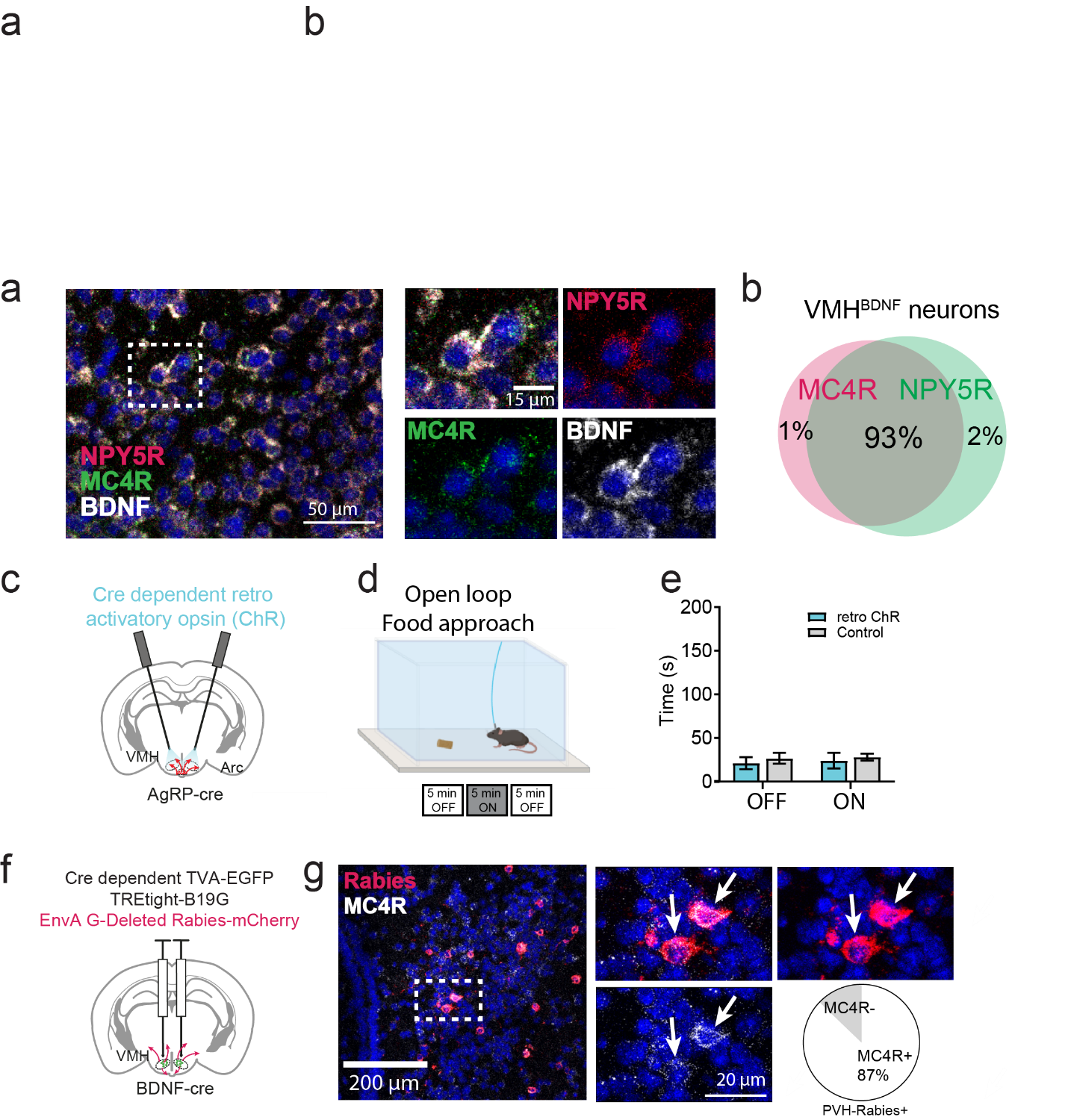

**Extended data figure 6**

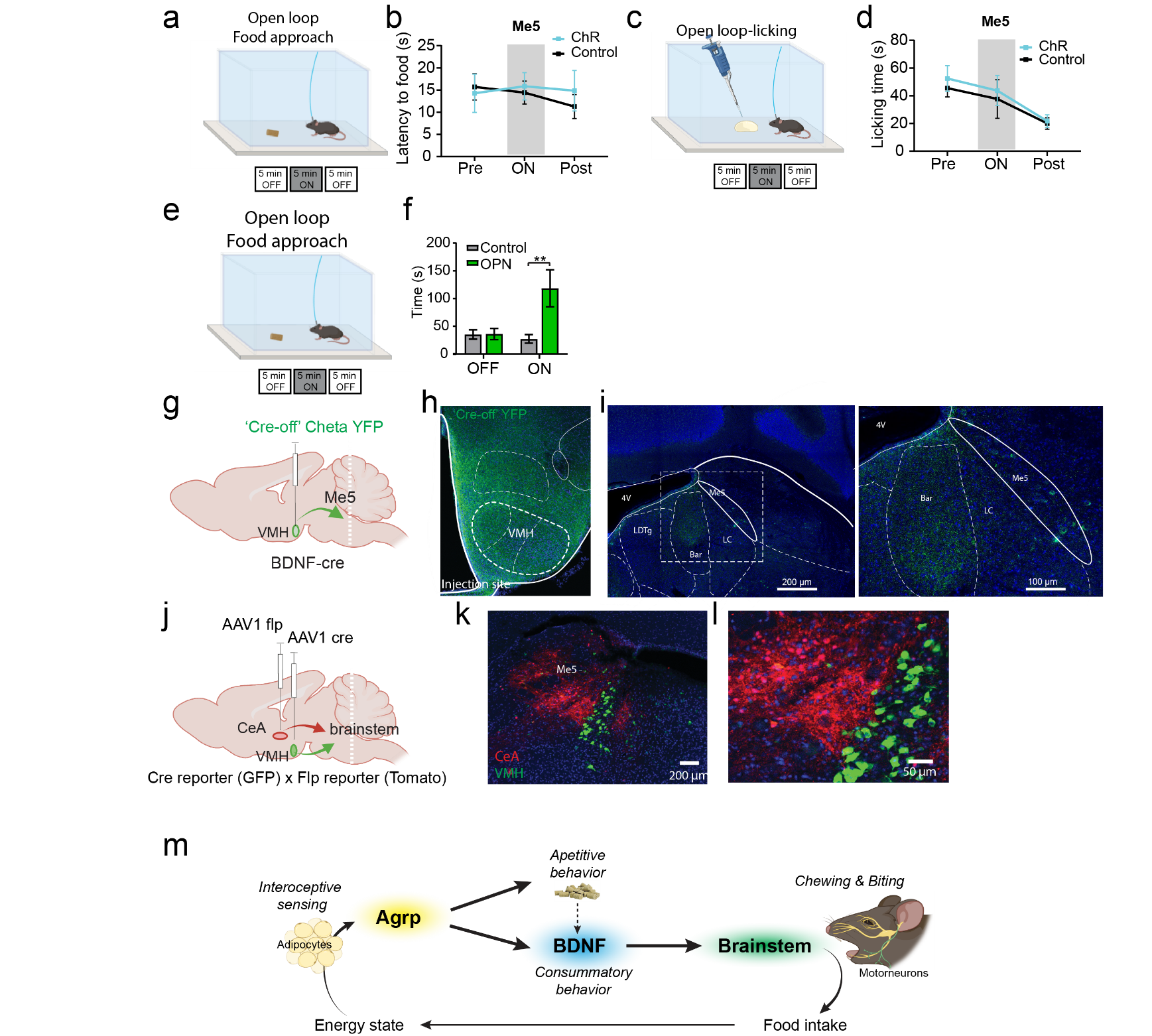
